# LIR-dependent LMX1A/LMX1B autophagy crosstalk shapes human midbrain dopaminergic neuronal resilience

**DOI:** 10.1101/636712

**Authors:** Natalia Jiménez-Moreno, Petros Stathakos, Zuriñe Antón, Deborah K. Shoemark, Richard B. Sessions, Ralph Witzgall, Maeve Caldwell, Jon D. Lane

**Author notes:** Corresponding author @jondlane.

## Abstract

The LIM homeodomain transcription factors LMX1A and LMX1B are essential mediators of midbrain dopaminergic neuronal (mDAN) differentiation and survival. Here we show that LMX1A and LMX1B are autophagy transcription factors in iPSC-derived human mDANs, each contributing to the expression of important autophagy genes including ULK1, ATG7, ATG16L1 and TFEB. Suppression of LMX1A and LMX1B in mDANs reduces basal autophagy, lowers mitochondrial respiration, and elevates mitochondrial ROS levels; meanwhile overexpression protects against rotenone poisoning in mDANs in vitro. Significantly, we show that LMX1A and LMX1B bind to multiple ATG8 proteins via LIR-type interactions, in a manner dependent on subcellular localisation and nutrient status: LMX1B interacts with LC3B in the nucleus under basal conditions via a C-terminal LIR, but binds to cytosolic LC3B and is degraded by autophagy during nutrient starvation, and LIR mutant LMX1B is unable to protect mDANs against rotenone. This establishes an LMX1A/LMX1B-autophagy regulatory nexus that helps explain the protective roles of these transcription factors in the adult midbrain, thus having implications for our understanding of mDAN decline in PD.

## INTRODUCTION

Parkinson’s disease (PD) is caused by the loss of midbrain dopaminergic neurons (mDANs) and the corresponding disruption of the nigrostriatal dopaminergic pathway (Kalia & Lang, 2015, Surmeier, Obeso et al., 2017). At the cellular level, the hallmarks of PD include intracellular accumulation of α-synuclein-enriched Lewy bodies, mitochondrial dysfunction and increased neuronal oxidative stress, and failing autophagy cytoplasmic quality control (Karabiyik, Lee et al., 2017, Surmeier et al., 2017). Autophagy protects cells through the delivery of toxic, damaged or redundant cytoplasmic cargo to the lysosome for degradation and recycling. In macroautophagy (henceforth referred to as “autophagy”), a novel organelle—the autophagosome—is assembled through the sequential actions of the products of conserved autophagy-related genes (ATG), and this organelle sequesters cargo in a selective or non-selective fashion for trafficking and fusion with late endosomes/lysosomes to form a degradative endolysosomal compartment (Lamb, Yoshimori et al., 2013). mDANs are particularly sensitive to autophagy deficits (Friedman, Lachenmayer et al., 2012, Karabiyik et al., 2017, Kett, Stiller et al., 2015, Sato, Uchihara et al., 2018), and correspondingly, upregulation of the autophagy and/or autolysosomal systems can protect against α-synuclein toxicity in mDANs in vivo (Decressac, Mattsson et al., 2013). Thus, understanding how autophagy responses are coordinated in mDANs is a key objective.

Neural precursor fate in the developing midbrain is determined by the precise spatiotemporal control of neuronal gene expression, coordinated by interdependent networks transcription factors (Doucet-Beaupre, Ang et al., 2015, Hegarty, Sullivan et al., 2013). Here, the LIM homeodomain transcription factors, LMX1A and LMX1B, stand out as essential determinants of mDAN differentiation and maintenance (Doucet-Beaupre et al., 2015). Sharing high amino acid homology (homeobox: 100%; LimA domain: 67%; LimB domain: 83% (Doucet-Beaupre et al., 2015)), LMX1A/LMX1B expression is spatiotemporally distinct: LMX1A is expressed early during differentiation, and regulates neurogenesis and neural fate in the ventral mesencephalic floor plate—consequently, mDANs are depleted (<50%) in the ventral midbrains of *Lmx1a* null and *dreher* mutant mice (Deng, Andersson et al., 2011, Yan, Levesque et al., 2011); meanwhile LMX1B, which has additional roles in dorsoventral patterning and 5-HT neuronal differentiation, indirectly guides mDAN differentiation via the isthmic organiser—a transient centre that controls midbrain-hindbrain regional identity through the regulated secretion of FGF8 and Wnt1 (Doucet-Beaupre et al., 2015). LMX1B also contributes to limb and kidney development (Doucet-Beaupre et al., 2015), with mutations in LMX1B being causative for nail-patella syndrome in humans (Harita, Kitanaka et al., 2016). *Lmx1a* and *Lmx1b* can partially compensate for one another in the mouse (Doucet-Beaupre et al., 2015, Yan et al., 2011), suggesting a degree of functional redundancy. Importantly, conditional knockout mouse models have unequivocally demonstrated the need for sustained *Lmx1a*/*Lmx1b* expression to support mDANs in the developed midbrain where *Lmx1a*/*Lmx1b* ablation triggered mDAN decline linked and neuropathological (e.g. α-synuclein-positive and distended axonal terminals) and behavioural abnormalities consistent with PD (Doucet-Beaupre, Gilbert et al., 2016, Laguna, Schintu et al., 2015). With this in mind, it is noteworthy that *Lmx1a* expression declines with age in the mouse brain (Doucet-Beaupre et al., 2016, Laguna et al., 2015), while LMX1B levels have been reported to inversely correlate with disease progression in the brains of PD patients (Laguna et al., 2015, Xia, Zhang et al., 2016). In addition, LMX1A/LMX1B polymorphisms have been linked (albeit weakly) to PD (Bergman, Hakansson et al., 2009). Thus, boosting and/or maintaining LMX1A/LMX1B levels in the adult brain may be of therapeutic benefit in PD.

Long-term and tissue tissue-specific autophagy responses are controlled at the transcriptional level by diverse families of transcription factors (Fullgrabe, Ghislat et al., 2016, Fullgrabe, Klionsky et al., 2014), but the transcriptional pathways that regulate autophagy gene expression in human mDANs remain elusive. Significantly, in the conditional *Lmx1a*/*Lmx1b* knockout mice, post-mitotic mDAN functional decline was associated with dysregulated autophagy, reduced mitochondrial function, and elevated mitochondrial oxidative stress (Doucet-Beaupre et al., 2016, Laguna et al., 2015). This suggests that LMX1A/LMX1B contribute to the expression of autophagy and mitochondrial quality control genes in mDANs. Using human iPSC-derived mDANs and HEK293T cells, we demonstrate here that LMX1A and LMX1B are indeed autophagy transcription factors that provide protection against PD-associated cellular stress in vitro. Importantly, both LMX1A and LMX1B bind to autophagy ATG8 family members in vitro, with binding of LMX1B to LC3B being dependent on a conserved LIR motif (^309^YTPL^312^). Interestingly, the LMX1B-LC3B interaction is restricted to the nucleus under full nutrient conditions, but shifts to the cytosol during starvation, leading to autophagy-dependent LMX1B turnover. We suggest that interplay between LMX1B and the core autophagy machinery influences its stability and transcriptional influence with implications for mDAN function and survival in the PD brain.

## RESULTS

### LMX1A and LMX1B are autophagy transcription factors in human mDANs

LMX1A and LMX1B emerged as distinct paralogs alongside the development of more complex chordate brain architecture (Holland, 2015) (**Appendix Fig S1A, B)**, but these transcription factors have spatiotemporally distinct expression patterns that are key to their roles during mDAN differentiation (Deng et al., 2011, Doucet-Beaupre et al., 2015, Yan et al., 2011). Given the possible link between LMX1A/LMX1B and the autophagy system, and its importance during mDAN maintenance/protection (Laguna et al., 2015), we used bioinformatics to search for LMX1A/LMX1B-targeted A/T-rich FLAT elements in autophagy gene promoter regions to provide evidence for LMX1A/LMX1B-mediated autophagy transcriptional control in humans (see **Materials & Methods**). As validation, FLAT elements were detected in known LMX1A/LMX1B transcriptional target promoters, including COL4A3 and IFNB1—both implicated in joint and/or kidney maintenance (Morello, Zhou et al., 2001, Rascle, Neumann et al., 2009)—and the mDAN control genes, NURR1, PITX3 and TH (Levesque & Doucet-Beaupre, 2013) (**Table EV1**). FLAT elements were also detected in the promoter of nuclear-encoded mitochondrial NDUFA2, but were also found in COX1 and NDUFV1 (LMX1B only) whose expression was not significantly altered following LMX1A/LMX1B combinatorial ablation in the Doucet-Beaupre et al. study (Doucet-Beaupre et al., 2016) (**Table EV1**). Importantly, putative FLAT elements were identified in genes controlling autophagy initiation (e.g. ULK1/2; ATG13; ATG14; WIPI2; UVRAG) and autophagosome expansion (e.g. ATG3; ATG5; ATG7; ATG10; ATG16L1) (**Table EV1**). These were also detected in the promoters of autophagy receptors (e.g. OPTN; NDP52; TAX1BP1; but not p62), and in both PINK1 and Parkin (**Table EV1**). The autophagy-related transcription factors TFEB and ZKSCAN3 also harboured putative FLAT elements for both LMX1A and LMX1B, while the hypoxia-responsive transcription factors NRF1 and NRF2 were positive for LMX1B only (**Table EV1**). This places LMX1A and LMX1B in the context of a wider cytoplasmic quality control transcriptional control network with the potential to protect against cellular stresses (Fullgrabe et al., 2016, Fullgrabe et al., 2014). Notably, co-isolation of tagged LMX1A and LMX1B overexpressed in HEK293T cells suggests that they have the potential to form heterodimers (**Appendix Fig S1C**), although the significance of this is as yet unclear.

Guided by the bioinformatics data, we selected a panel of candidate genes for LMX1B ChIP analysis in human HEK293T kidney cells that express endogenous LMX1B (Burghardt, Kastner et al., 2013). Using an arbitrary 2-fold cut-off for targets of interest, promoter occupancy was confirmed for several core autophagy genes, including ULK1, ATG3, ATG16L1, UVRAG, as well as the autolysosomal transcription factor TFEB and the receptor and/or mitophagy genes NDP52, OPTN and PINK1 (**Fig 1A**). The mDAN differentiation genes ABRA, NURR1 and PITX3 were also identified in LMX1B ChIP in HEK293T cells (**Fig 1A**). To provide experimental support for its role as a human autophagy transcription factor, we measured candidate autophagy gene expression by qRT-PCR in HEK293T cells depleted for LMX1B using siRNA (**Fig 1B**) or shRNA (**Fig 1C**). Expression of several autophagy genes including ULK1, ATG3, ATG7, ATG16L1, UVRAG, TFEB was significantly reduced following LMX1B suppression, as was expression of the selective autophagy/mitophagy receptors NDP52 and OPTN, and the mitophagy regulator and early-onset PD gene, PINK1 (**Fig 1B, C**). ATG3 was significantly reduced following treatment with siLMX1B, but not when using shLMX1B (**Fig 1B, C**), perhaps due to the relative efficiencies of LMX1B suppression. p62 was not affected by LMX1B suppression, confirming bioinformatics predictions, but neither was ATG5 despite having been identified as potentially possessing FLAT element(s) in the bioinformatics dataset (**Table EV1**). Importantly, the expression of several autophagy genes could be rescued in siLMX1B-treated HEK293T cells by overexpression of siRNA-resistant LMX1B (**Appendix Fig S1D**). These findings were corroborated for several genes at the level of protein expression, including ATG3 and ATG7 (significantly reduced), although expression of ATG16L1 was not affected (**Fig 1D**). Indicative of reduced autophagy potential in siLMX1B-treated HEK293T cells, basal levels of lipidated LC3II were significantly lower (**Fig 1D**), and this was confirmed by analysis of LC3B, and WIPI2 puncta numbers in HEK293T cells (**Fig 1E-H**). p62 puncta were also found to decrease following LMX1B siRNA (**Fig 1I**), perhaps suggesting that p62 association with puncate (ubiquitylated) cargo is suppressed. Together, these data demonstrate that LMX1B contributes to the regulation of autophagy gene expression in HEK293T cells, and that LMX1B suppression dampens the basal/housekeeping autophagy response in this setting.

**Figure 1.**
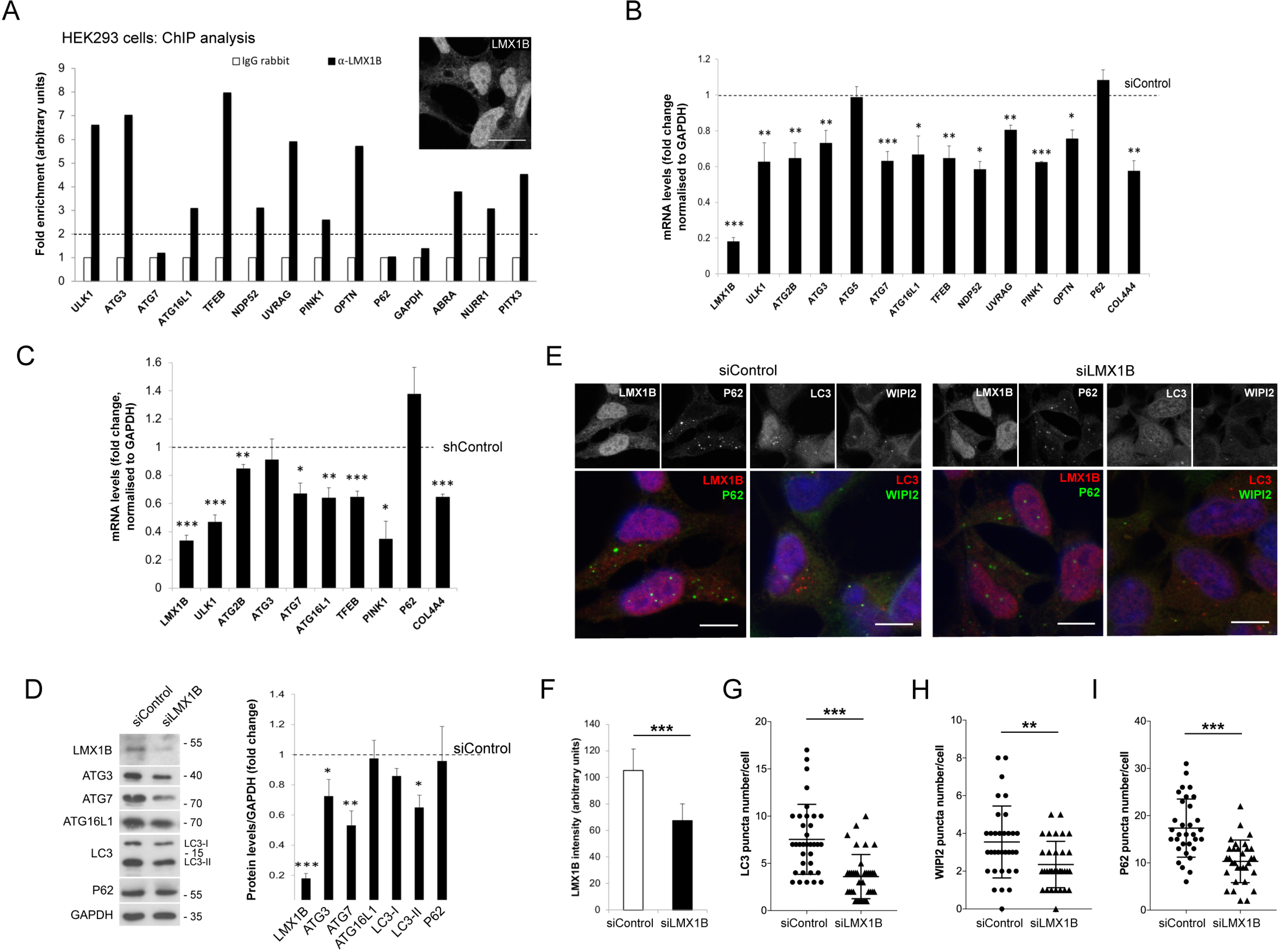
**LMX1B is an autophagy transcription factor in HEK293 cells.** (A) ChIP qRT-PCR analysis of LMX1B promoter occupancy in HEK293T cells. This experiment was repeated three times with similar results. Inset shows nuclear anti-LMX1B staining in HEK293T cells. Bar = 20µm. (B, C) qRT-PCR analysis of selected genes in HEK293T cells transfected with (B) LMX1B siRNA smartpool or non-targeting siControl, or (C) shLMX1B or non-targeting shControl. mRNA levels were normalized to GAPDH. Mean ± SEM (n = 3); student’s t test: *p<0.05, ** p<0.01 and *** p<0.001 vs. si/shControl. (D) LMX1B siRNA reduces expression of key autophagy proteins. Left, example immunoblots; right, quantitation relative to GAPDH levels. Mean ± SEM (n = 3); student’s t test: *p<0.05, ** p<0.01 and *** p<0.001 vs. siControl. (E) GFP-LC3B HEK293T cells transfected with non-targeting siControl (left) or siLMX1B (right), stained with anti-LMX1B, anti-WIPI2 or anti-p62 antibodies. GFP-LC3B and LMX1B are depicted in red; WIPI2 and p62 are depicted in green. Cells were counterstained with DAPI (blue). Bar = 10µm. (F-I) LMX1B intensity (F), and puncta counts for GFP-LC3B (G), WIPI2 (H), and p62 (I) in siControl and siLMX1B-treated HEK293T cells. Mean ± SD of >35 cells from three independent experiments; student’s t test: ** p<0.01 and *** p<0.001 vs. siControl.

To confirm their potential as autophagy transcription factors in the human midbrain, we prepared mDANs from the NAS2 (normal α-synuclein) iPSC line (Devine, Ryten et al., 2011), using an optimised monolayer protocol that generates high numbers of mDANs expressing midbrain, dopaminergic markers including TH, LMX1A, LMX1B, and FOXA2 (Stathakos, Jimenez-Moreno et al., 2019) (see **Materials & Methods**). For these experiments, our cultures comprised ∼75% neurons, of which ∼50% expressed TH (**Fig 2A**). Of those TH-positive neurons, ∼80% expressed LMX1A and/or LMX1B (**Fig 2A**). Expression of LMX1A and LMX1B peaked and remained high from D20 of mDAN differentiation, correlating with the expression of neuronal βIII tubulin (TUJ1) and the dopaminergic transcription factor, NURR1 (**Fig 2B-E**). Based on an arbitrary 2-fold cut-off, ChIP analysis of iPSC mDAN cultures using anti-LMX1A and anti-LMX1B antibodies indicated promoter occupancy for the autophagy genes ATG7 (for LMX1A/LMX1B), ATG16L1 (LMX1A only) and ULK1 (LMX1B only), and the adaptors NDP52 and OPTN (**Fig 2F, G**). Dopaminergic neuronal control genes NURR1 and PITX3 were also positive for both LMX1A and LMX1B, with ABRA being positive for LMX1B only (**Fig 2G**), as expected (Burghardt et al., 2013) (**Table EV1**). To test the impact of LMX1A/LMX1B suppression in human mDANs in vitro, we generated lentiviruses expressing GFP (synapsin promoter) with shRNA (U6 promoter) (**Fig 2H**). 6-days post viral transduction in D>30 cultures, LMX1A and LMX1B expression in mDANs was significantly reduced (**Fig 2H**), with suppression of either gene significantly reducing the other (**Fig 2I**). Importantly, shRNA suppression of either LMX1A or LMX1B significantly reduced expression of ULK1, ATG2B, ATG7, ATG16L1, NDP52, OPTN, PINK1 and TFEB in mDANs, but ATG3 and p62 were not affected (**Fig 2I**). Expression of the dopaminergic neuronal genes NURR1 and PITX3 were significantly reduced in both conditions as expected (Doucet-Beaupre et al., 2015, Nakatani, Kumai et al., 2010), but TH and TUJ1 expression levels were not altered within this time-frame (**Fig 2I**). Expression of MSX1 was significantly reduced only in the LMX1A suppressed cells (**Fig 2I**), consistent with previous reports that MSX1 is controlled by LMX1A in the midbrain (Andersson, Tryggvason et al., 2006, Cai, Donaldson et al., 2009), but as LMX1B suppression concomitantly reduced LMX1A levels (**Fig 2I**), this implied that residual LMX1A levels must be sufficient to maintain MSX1 levels in this case.

**Figure 2.**
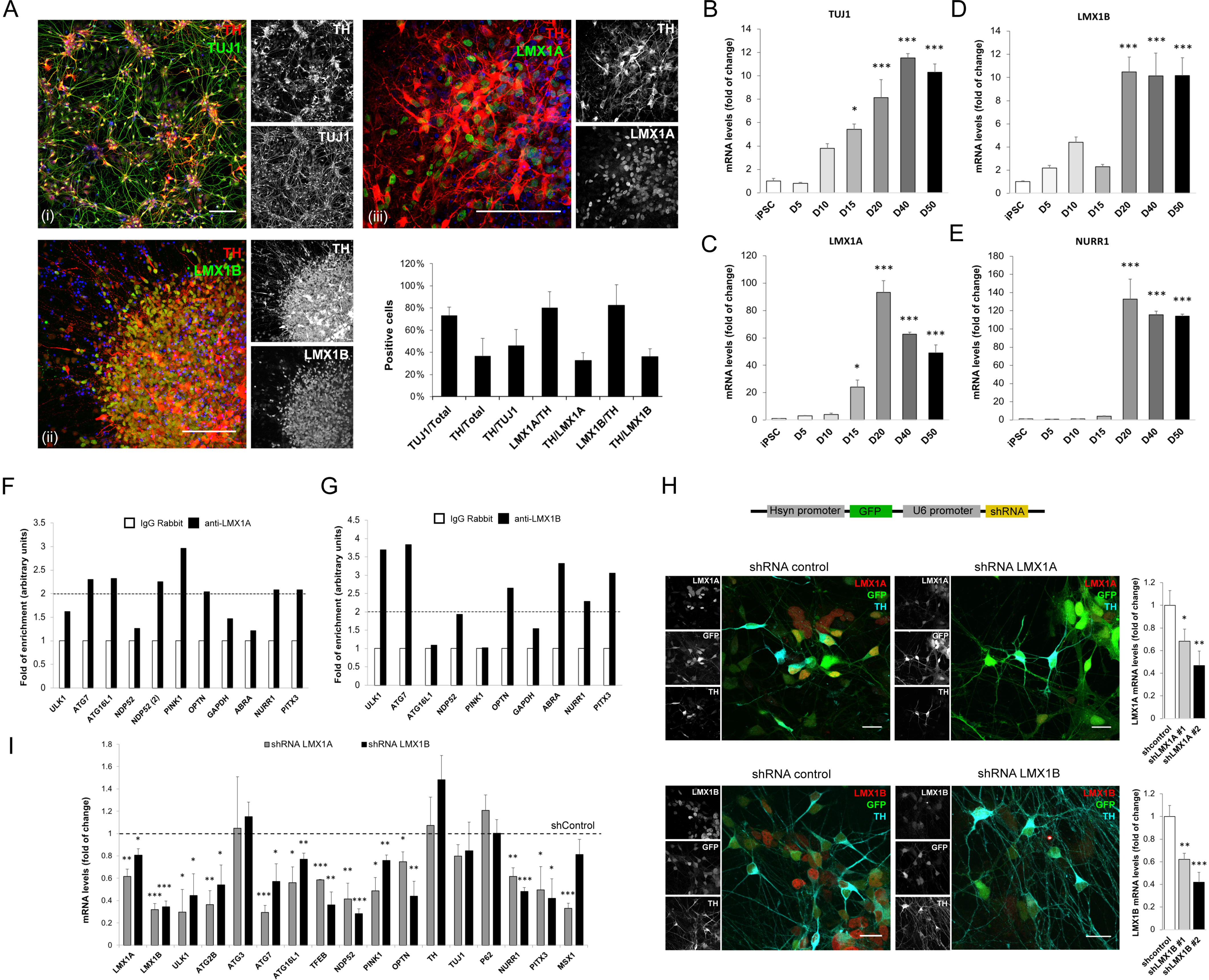
LMX1B is an autophagy transcription factor in iPSC-derived human mDANs. (A) Immunofluorescence images and quantitation of LMX1A/LMX1B and dopaminergic marker expression in human iPSC-derived mDANs (imaged at: [i] D40; [ii, iii] D20). Cells were stained for anti-TH (red) and either anti-TUJ1 (green), anti-LMX1A (green) or anti-LMX1B (green). Cells were counterstained with DAPI (blue). Bar = 100µm. Data show counts of cells positive for the marker combinations shown (mean ± SD; n=3). (B-E) qRT-PCR analysis of (B) TUJ1, (C) LMX1A, (D) LMX1B, and (E) NURR1 differentiated at days 5, 10, 15, 20, 40 and 50 (or iPSC control). Levels were normalized to GAPDH. Mean ± SEM (n = 3) of mDAN cultures plated from a single neuralization; one-way ANOVA followed by a Dunnett’s multiple comparisons test. *p<0.05, ** p<0.01 and *** p<0.001 vs. iPSCs. (F, G) ChIP qRT-PCR analysis of (F) LMX1A and (G) LMX1B promoter occupancy in human mDANs. (H) shRNA suppression of LMX1A/LMX1B expression in iPSC-derived mDANs. A schematic representation of the pRRL plasmid construct expressing GFP under human synapsin promoter (hsyn) and shRNA for LMX1A or LMX1B (or a non-target shRNA control) under the U6 promoter is shown to the top. Example fields of mDANs stained for anti-LMX1A (top, red) or anti-LMX1B (bottom, red) and anti-TH (cyan) after transduction with hsyn-GFP-U6-shControl, hsyn-GFP-U6-shLMX1A or hsyn-GFP-U6-shLMX1B (green). To the right, LMX1A and LMX1B qRT-PCR quantitation after viral transduction at D30-45 (normalized to GAPDH) of two different shRNAs for LMX1A and LMX1B. Mean ± SEM (n=3); student’s t test: *p<0.05, ** p<0.01 and *** p<0.001 vs. shRNA control. Bars = 20µm. (I) qRT-PCR analysis of candidate gene expression in iPSC-derived mDANs (D30-45) following hsyn-GFP-U6-shControl/shLMX1A/shLMX1B lentiviral transduction (normalized to GAPDH). Means ± SEM (n=3); student’s t test: *p<0.05, ** p<0.01 and *** p<0.001 vs. shRNA control.

### LMX1A and LMX1B regulate autophagy and protect human iPSC-derived mDANs in vitro

LMX1A and LMX1B suppression significantly reduced the numbers of WIPI2-positive autophagosome assembly sites in the cell bodies of mature (D>30) TH- and GFP-positive mDANs consitent with a dampened basal autophagy response (**Fig 3A**). Indicative of increased mitochondrial stress, human mDANs suppressed for LMX1A or LMX1B showed evidence of elevated mitochondrial ROS (MitoSox; **Fig 3B**), associated with reduced basal oxygen consumption rate, maximal respiration, and spare capacity (**Fig 3C, D**), consistent with declining mitochondrial fitness (as seen in isolated mouse mDANs lacking LMX1A/LMX1B (Doucet-Beaupre et al., 2016)). In human iPSC-derived mDANs, the timing of LMX1A/LMX1B suppression was critical: neurite extension kinetics following plating of mDANs were not affected by LMX1A/LMX1B shRNA at D>35 (**Appendix Fig S2A**), but infection during the rapid LMX1A/LMX1B expression phase (i.e. D<17) negatively impacted neurite length following a ∼12 hour lag (**Appendix Fig S2B**). This was associated with marked changes in expression of dopaminergic genes (TUJ1, TH, NURR1, MSX1) in D<17 cultures suppressed for LMX1A, with LMX1B suppression associated with compensatory LMX1A elevation and enhanced NURR1 expression (**Appendix Fig S2C**). Despite this, population cell death levels measured by caspase-3 assay remained similar to control shRNA-treated mDANs (**Fig 3E**). Thus, mature mDANs can cope well in the absence of either transcription factor in vitro, at least within this short time-frame.

**Figure 3.**
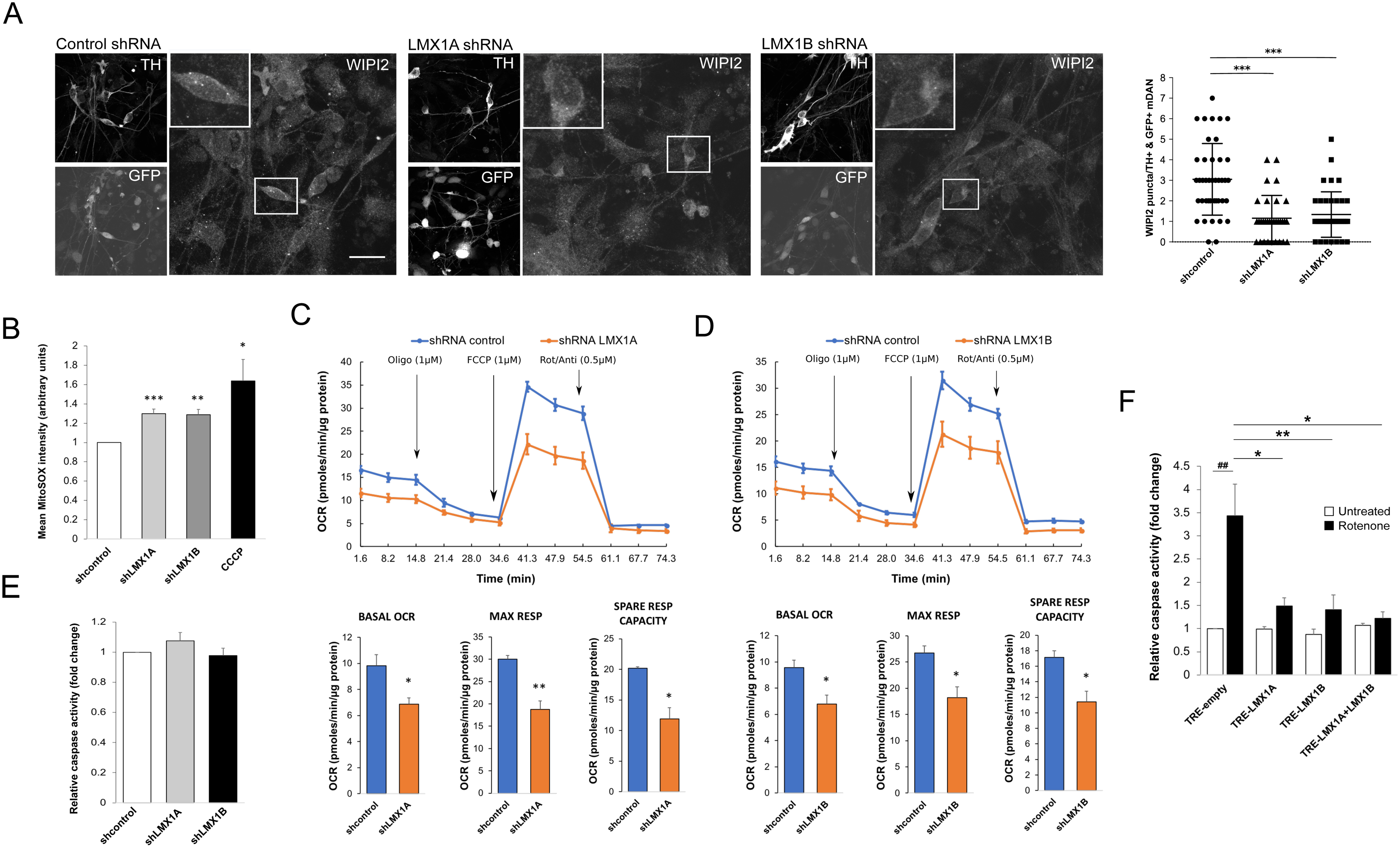
LMX1A/LMX1B control basal autophagy and mitochondrial function to protect mDANs. (A) Representative immunofluorescence images and quantitation of basal WIPI2 puncta in TH/GFP positive iPSC-derived mDAN cell bodies (D30-45) following hsyn-GFP-U6-shControl/shLMX1A/shLMX1B lentiviral transduction. Mean ± SD (>30 neurons/condition; three independent experiments); student’s t test: *** p<0.001 vs. shRNA control. Bar = 20µm. (B) MitoSOX intensity in TH-positive iPSC-derived mDANs transduced with hsyn-GFP-U6-shControl/shLMX1A/shLMX1B (D30-50). Cells were treated with CCCP (10µM, 4h) as a positive control. Mean ± SEM of 3 experiments, >30 neurons per condition; student’s t test: *p<0.05, ** p<0.01 and *** p<0.001 vs. shRNA control. (C, D) Mitochondrial respiration profiles in iPSC-derived mDAN cultures (D30-50) transduced with (C) hsyn-GFP-U6-shRNA LMX1A or (D) hsyn-GFP-U6-shRNA LMX1B lentiviruses. Pairwise comparisons were performed against non-targeting shRNA control. Oxygen consumption rate (OCR) was normalized to protein levels after sequential exposure to oligomycin (1µM), FCCP (1µM) and Rotenone/Antimycin (0.5µM) as indicated. Mean ± SEM (n = 3); student’s t test: *p<0.05 and **p<0.01 vs. shRNA control. (E) Basal mDAN cell death analysis following hsyn-GFP-U6-shControl/shLMX1A/shLMX1B lentiviral transduction (D25-50 mDANs). Fluorometric caspase assay normalized to total protein. Mean ± SEM (n=3) (not significant). (F) Rotenone-induced cell death in iPSC-derived mDANs (D25-50) transduced with TRE-empty (control), TRE-LMX1A, and/or TRE-LMX1B; expression induced by doxycycline (500ng/mL, 3 days). mDANs were treated with rotenone (15µM) for 24h. Caspase activity measured relative normalized to total protein. Mean ± SEM (n=3); one-way ANOVA followed by a Tukey’s multiple comparison post-hoc test: *p<0.05 and **p<0.01 vs rotenone-treated and ^##^p<0.01 untreated vs rotenone-treated.

Overall, these tests show that LMX1A or LMX1B suppression is well tolerated in the short-term in mature (D>30) iPSC-derived mDANs in vitro, but correlates with reduced basal autophagy, impaired mitochondrial function, and increased mitochondrial ROS. Crucially, using a doxycycline inducible LMX1A/LMX1B lentiviral system, we found that overexpression of LMX1B alone or in combination with LMX1A increased the expression of several key autophagy-associated genes in human mDANs (notably, ULK1, ATG3, TFEB; **Appendix Fig S2D**). LMX1A overexpression by itself did not induce autophagy gene transcription (**Appendix Fig S2D**), as also reported in the mouse (Laguna et al., 2015). LMX1B overexpression concomitantly increased LMX1A levels, but the converse was not the case (**Appendix Fig S2D**). The significance of these observations became clear when we measured stress-induced mDAN cell death by treatment with the mitochondrial poison, rotenone (a complex I inhibitor widely used as a PD-inducing model; e.g (Betarbet, Sherer et al., 2000)). In our mDAN cultures, induced LMX1A and/or LMX1B overexpression significantly reduced rotenone-induced cell death compared to the empty vector control (**Fig 3F**). Interestingly, LMX1A overexpression was cytoprotective in this assay (**Fig 3F**), despite this transcription factor not independently driving expression of any of the autophagy-related genes that we tested (**Appendix Fig S2D**), perhaps suggesting involvement of other protective LMX1A transcriptional targets.

### Autophagy-dependent turnover of the LMX1B protein

The apparent correlation between LMX1B levels and mDAN protection from PD-related mitochondrial stress, and the evidence that reduced LMX1B expression impairs autophagy responses in mDANs ((Laguna et al., 2015); this study) and correlates with ageing and PD progression (Doucet-Beaupre et al., 2016, Laguna et al., 2015), prompted us to monitor LMX1B turnover. Degradation of LMX1B-FLAG in HEK293T cells during cycloheximide treatment (16 hours) was blocked by Bafilomycin A1 (BafA1; lysosomal inhibitor) and by MG132 (proteasome inhibitor) (**Fig 4A**), suggesting involvement of both autolysosomal and proteasomal degradation pathways. Significantly, we found that LMX1B-FLAG turnover in HEK293T cells was autophagy-dependent during nutrient starvation (**Fig 4B**). As the distribution of key transcription factors such as TFEB, ZSCAN3 and FOXO alters during stress (Fullgrabe et al., 2016), we used cell fractionation and imaging to monitor the localisation of LMX1B-FLAG in HEK293T cells under nutrient stress (**Figs 4C, D**). Immunoblotting revealed that the nuclear LMX1B-FLAG fraction increased significantly, while the cytosolic LMX1B-FLAG component was correspondingly reduced, during starvation (2 hours), suggesting net movement from cytoplasm to nucleus in response to nutrient stress (**Fig 4C**). Observed by microscopy, LMX1B-FLAG localisation in HEK293T cells in full media and during acute (2 hours) starvation was predominantly nuclear, with small numbers of additional cytosolic LMX1B-FLAG foci observed in some cells (**Fig 4D**). These foci remained distinct from autophagic membranes labelled with GFP-LC3B (**Fig 4D**). Strikingly, following extended starvation (6 hours), LMX1B-FLAG translocated to the cytosol where it accumulated in abundant cytosolic puncta that strongly co-localised with GFP-LC3B (**Fig 4D**). Thus, in common with TFEB whose nucleus-to-cytoplasm shuttling is regulated by nutrient availability (Puertollano, Ferguson et al., 2018), LMX1B accumulates in the nucleus during acute starvation; however, LMX1B exits and is degraded in the cytosol during extended periods of nutrient starvation. This suggests a possible feedback mechanism where early nutrient starvation triggers LMX1B translocation to the nucleus where it can boost the autophagy response, but during prolonged stress, LMX1B is subsequently re-exported to be targeted by the autophagy pathway to limit the duration of this protection.

**Figure 4.**
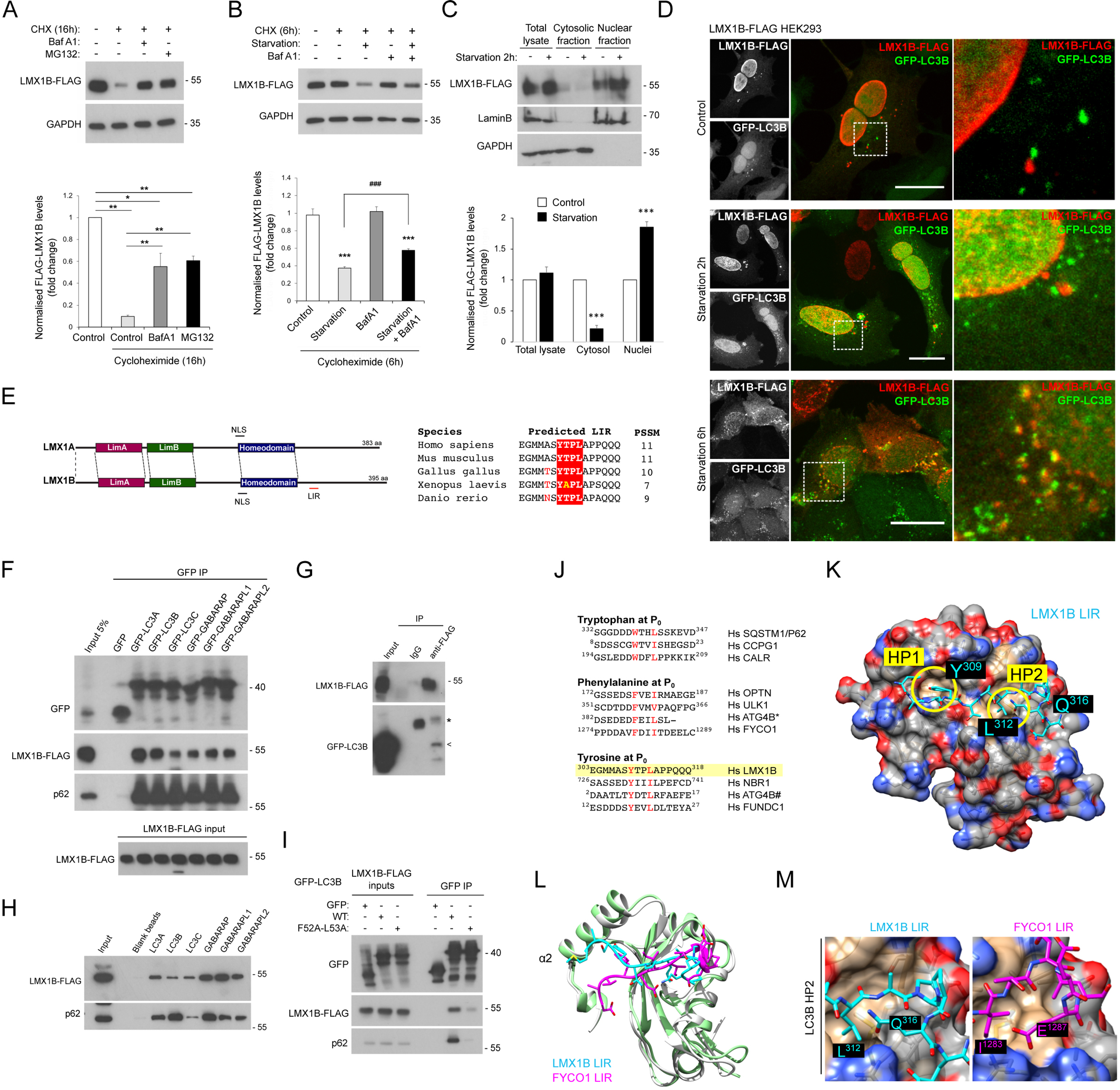
LMX1B interacts with ATG8s and is degraded during nutrient starvation. (A) LMX1B-FLAG turnover in HEK293T cells treated with cycloheximide (CHX, 50µg/mL) for 16h in the absence or presence of MG132 (10µM) or BafA1 (20nM). Mean ± SEM (n = 3); one-way ANOVA followed by Tukey’s multiple comparison post-hoc test: *p<0.05 and ** p<0.01. (B) LMX1B-FLAG degradation during starvation (6h) in the presence of CHX ± BafA1. Mean ± SEM (n = 3); one-way ANOVA followed by Dunnett’s multiple comparisons test (*p<0.05 and *** p<0.001 vs. control), and student’s t test: ^###^p<0.001 starvation vs. starvation + BafA1. (C) Immunoblot and densitometry of cytosolic and nuclear fractions from HEK293T cells expressing LMX1B-FLAG with or without 2h nutrient starvation. Lamin B was used to normalize the nuclear fractions and GAPDH for the total and cytosolic fractions. Mean ± SEM (n = 3); student’s t test: *** p<0.001 vs. untreated controls. (D) LMX1B relocates to cytosolic puncta that co-label with LC3B during prolonged nutrient starvation. Representative immunofluorescence images of HEK293T cells co-transfected with GFP-LC3B (green) and LMX1B-FLAG, fixed and stained with anti-FLAG antiserum (red) after 0h, 2h or 6h starvation. Bar = 20µm. (E) Domain schematic of human LMX1A and LMX1B showing predicted nuclear localisation sequences (NLS) and a possible LIR motif in LMX1B (left) (the same schematic is shown in **Appendix Fig S3A** depicting the location of a possible LIR in LMX1A). Alignment of the LMX1B LIR in different species (right). (F) GFP-TRAP co-precipitation of LMX1B-FLAG with GFP-ATG8 family members in HEK293T cells. 5% protein lysate from equivalent GFP-expressing cells is shown as “input”. (G) Anti-FLAG co-precipitation of GFP-LC3B from lysates of HEK293T cells stably expressing FLAG-LMX1B. A non-specific IgG control is included. Arrow indicates position of GFP-LC3B; * indicates the position of the antibody heavy chain. (H) In vitro pull-down of LMX1B-FLAG from lysates of HEK293T stably expressing FLAG-LMX1B using sepharose beads covalently attached to recombinant His-ATG8 family members. (I) GFP-TRAP co-precipitation of LMX1B-FLAG with wild-type of LIR docking mutant (F52A/L53A) GFP-LC3B in HEK293T cells 5% of protein lysate was used as control for protein expression (input). (J) Sequence alignments of LIR motifs in various human (Hs) proteins designated as tryptophan-, phenylalanine- and tyrosine-type LIRs (refers to the residue at P_0_ of the LIR). * C-terminal ATG4B LIR; # N-terminal ATG4B LIR. (K) LMX1B LIR overlayed upon a space-filling model of LC3B (5d94.pdb) in final simulation pose, showing docking of the key LIR residues at P_0_ and P_3_ within hydophobic pockets (HP) 1 and 2 of LC3B respectively. (L) Ribbon structure overlay of human LC3B (grey) in complex with the FYCO1 LIR (Magenta) (5d94.pdb) with the final model simulation pose of LC3B (green) and LMX1B LIR (cyan). Close alignment between FYCO1 and LMX1B is seen within the core LIR binding region. (M) Side-by-side comparison of the LMX1B (left) and FYCO1 (right) LIRs docked at HP2 of LC3B, (5d94.pdb) to show how LMX1B Q316 and FYCO1 E1287 fold back towards HP2 in both structures to stabilise LIR binding.

### LMX1B contains a LIR motif and bind multiple ATG8 proteins via LIR-type interactions

Analysis of the primary structure of LMX1B using the iLIR search tool (https://ilir.warwick.ac.uk) revealed the presence of a possible LIR motif distal to the C-terminal homeodomain (**Fig 4E**). LIR motifs are present in proteins that interact with ATG8 family members to facilitate their autophagy-mediated turnover, and in proteins that act as receptors for selective autophagy cargo and/or to contribute to the autophagosome assembly pathway (Lamark, Svenning et al., 2017). We therefore tested whether LMX1B interacted with ATG8 family members by GFP-TRAP immunoprecipitation of lysates of HEK293T cells co-transfected with human GFP-ATG8 and FLAG-LMX1B (**Fig 4F**). Immunoblots revealed that LMX1B-FLAG was enriched in all GFP-ATG8 fractions (as was p62; **Fig 4F**). GFP-LC3B was also detected in anti-FLAG immunprecipitates from lysates of HEK293T cells stably expressing LMX1B-FLAG (**Fig 4G**), and LMX1B-FLAG bound to ATG8 beads in vitro, with a possible preference for GABARAPs in this context (**Fig 4H**). Consistent with interactions being LIR-dependent, LIR docking site mutant GFP-LC3B (F52A/L53A) (Ichimura, Kumanomidou et al., 2008) co-precipitated very weakly with LMX1B-FLAG (**Fig 4I**). Interestingly, iLIR also identified a putative LIR motif in human LMX1A, and LMX1A co-precipitated with GFP-ATG8s in HEK293T cells in a LIR-dependent manner (**Appendix Fig S3A-C**), suggesting that both transcription factors interact with the autophagy system (although alanine mutagenesis of the putative LMX1A LIR motif (^290^YTAL^293^) did not alter binding to LC3B; **Appendix Fig S3D**).

Alignment of the predicted C-terminal LIR identfied in LMX1B (^309^YTPL^312^) against other published LIR motifs suggested that it conforms to the tyrosine-type group identified in LIR domain containing proteins including NBR1, ATG4B and FUNDC1, but with a relatively weak upstream acidic stretch (**Fig 4J**). Using Chimera (Pettersen, Goddard et al., 2004) we visualised the LC3B crystal structure bound to the FYCO1 LIR peptide (5d94.pdb) (Olsvik, Lamark et al., 2015), and used this structure as a template to guide the positioning of the LMX1B LIR and its flanking sequences. Throughout 4 repeat molecular dynamics simulations of the resulting assemblies (see **Materials & Methods**), the LMX1B LIR peptide remained in contact with LC3B (**Appendix Fig S4A**; **Movie EV1**), with critical Y309 and L312 anchors docked within LC3B hydrophobic pockets HP1 and HP2, respectively (**Fig 4K**). Recent data have highlighted the importance of residues outside of the core LIR in stabilising interactions with ATG8 proteins (Wirth, Zhang et al., 2019). In the case of the LMX1B LIR, differences were observed in the positioning of upstream residues relative to the FYCO1 crystallographic position, with the LMX1B LIR adopting a final pose that was displaced towards the α2 helix of the N-terminal arm of LC3B (Birgisdottir, Lamark et al., 2013) (**Fig 4L**). The FYCO1 LIR residues D1277 and D1281 form salt bridges between R10 and R70 of LC3B respectively. By contrast, the LMX1B LIR has no negatively charged residues in this region; rather A307 (i.e. at LIR X_-2_; **Fig 4J**) of the LMX1B LIR packs against L22 of LC3B. Downstream of the presumptive LIR, the first glutamine in the LMX1B triple Q stretch (i.e. at LIR X_7_), folded back in the simulation to adopt a simular configuration to that reported for the glutamic acid at LIR X_7_ in FYCO1 (**Fig 4M**), thus possibly stabilising the interaction. Interestingly, our proposed LMX1B-LC3B model is highly similar to the docking of the N-terminal tyrosine-type LIR of human ATG4B (see **Fig 4J**) seen in the lattice structure of the ATG4B-LC3B crystal (Satoo, Noda et al., 2009), both within the core LIRs, and in the manner in which the displaced upstream sequences engage with the LC3B α2 helix (**Appendix Fig S4B**).

### The LIR-dependent LMX1B-LC3B interaction is location and context dependent

At steady state, a large and relatively immobile pool of LC3 localises to the nucleus where it engages with PML bodies and nucleolar components via LIR-type interactions suggesting roles nuclear regulatory roles (He, Hu et al., 2014, Kraft, Manral et al., 2016). Notably, it has been demonstrated that cytosol-to-nuclear shuttling forms part of the normal LC3 itinerary, with only the nuclear released pool competent to undergo lipidation during autophagosome assembly (Huang, Xu et al., 2015). Strikingly, we found that the LMX1B-LC3B interaction occured exclusively in the nuclear compartment in LMX1B-FLAG HEK293T cells under full nutrient conditions (**Fig 5A**). Indeed, LMX1B binding to all ATG8 family members was restricted to the nucleus, unlike p62 which bound ATG8 family members overwhelmingly in the cytoplasm (**Appendix Fig S5**). To test whether nutrient availability influenced the location and apparent strength of LMX1B binding to LC3B, HEK293T cells co-expressing GFP-LC3B with LMX1B-FLAG were placed in starvation media for 2 hours, then cell fractions were subjected to GFP-TRAP immunoprecipitation (**Fig 5B**). This revealed a new interaction between LMX1B and GFP-LC3B in the cytosol in starved cells (**Fig 5B**). By contrast, cytosolic p62 binding to LC3B was not markedly influenced by starvation (**Fig 5B**). The steady state LMX1A interaction with GFP-LC3B in fed cells occurred primarily in the cytosol (**Appendix Fig S3E**), although this should be interpreted with some caution since LMX1A (and possible also its binding partners) are not normally expressed in HEK293T cells. In common with LMX1B, however, this interaction strengthened during starvation (**Appendix Fig S3F**), and LMX1A expressed in HEK293T cells was turned over by both lysosomal and proteasomal routes (**Appendix Fig S3G**).

**Figure 5.**
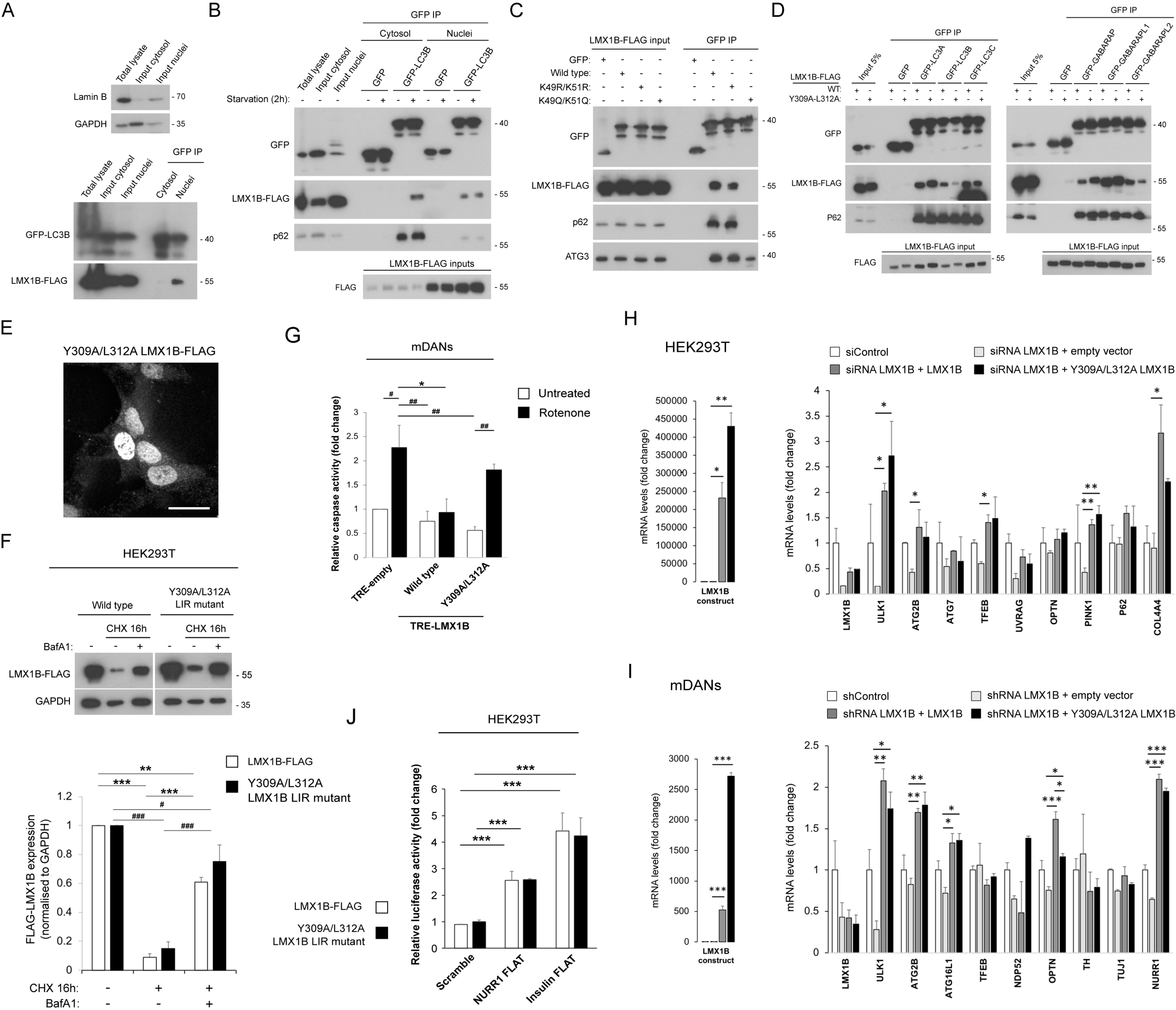
**LMX1B binds LC3B in the nucleus to promote mDAN survival.** (A) GFP-TRAP immunoprecipitation of nuclear and cytosolic fractions from lysates of HEK293T cells co-expressing GFP-LC3B and LMX1B-FLAG. The LMX1B-LC3B interaction occurs exclusively in the nucleus under basal conditions. (B) GFP-TRAP immunoprecipitation of nuclear and cytosolic fractions from lysates of HEK293T cells co-expressing GFP-LC3B and LMX1B-FLAG. Comparisons of pull-downs in full nutrients or following 2h starvation. (C) GFP-TRAP immunoprecipitation of LMX1B-FLAG in HEK293T cells co-expressing wild-type, acetylation-deficient (K49R/K51R) and acetylation mimic (K49Q/K51Q) GFP-LC3B. (D) GFP-trap immunoprecipitates of wild-type or LIR mutant (Y309A/L312A) LMX1B-FLAG in lysates of HEK293T co-expressing GFP-ATG8 family members. 5% of protein lysate from equivalent GFP-expressing cells is shown as a representative of “input”. (E) Immunofluorescence staining of LIR mutant (Y309A/L312A) LMX1B-FLAG expressed in HEK293T cells. Bar = 20µm. (F) LMX1B turnover in HEK293T expressing wild-type or Y309A/L312A LMX1B-FLAG. Cells were treated with CHX (50µg/mL; 16h) in the absence or presence BafA1 (20nM). Representative blot (top) and quantitation relative to GAPDH (bottom). Mean ± SEM (n = 3); one-way ANOVA followed by Tukey’s multiple comparison post-hoc test: **p<0.01 and ***p<0.001 vs LMX1B-FLAG and ^#^p<0.05 and ^###^p<0.001 vs Y309A/L312A LMX1B-FLAG. (F) Rotenone-induced cell death (15µM; 24h) of iPSC-derived mDAN cultures (D30-50) transduced with TRE-empty (control), TRE-LMX1B, or TRE-Y309A/L312A LMX1B (expression induced with doxycycline: 500ng/mL; 3d). Active caspase fluorescence relative to total protein levels was normalized to TRE-empty untreated values. Mean ± SEM (n=3); one-way ANOVA followed by Tukey’s multiple comparison post-hoc test: *p<0.05 vs rotenone-treated and ^#^p<0.05 and ^##^p<0.01 untreated vs rotenone-treated cells. (G) HEK293T cells were double co-transfected with 50nM smartpool LMX1B siRNA and codon optimised, siRNA-resistant wild-type or Y309A/L312A LIR mutant LMX1B (or empty vector control). Mean ± SD (n = 3 replicate wells); one-way ANOVA followed by Tukey’s multiple comparison post-hoc test comparing the three different siRNA conditions: *p<0.05 and ** p<0.01. (H) iPSC-derived mDANs (D∼50) were transduced with the hsyn-GFP-U6-shLMX1B and LMX1B levels were rescued with codon optimised, shRNA-resistant wild-type or Y309A/L312A LIR mutant TRE-GFP-LMX1B (or empty vector as a control). Mean ± SD (n = 3 wells from a single neuralisation); one-way ANOVA followed by a Tukey’s multiple comparison post-hoc test comparing the three different siRNA conditions: *p<0.05, ** p<0.01 and *** p<0.001. (I) Induced expression of NURR1 and pro-insulin FLAT promoter-driven luciferase (LightSwitch™) by wild-type or LIR mutant (Y309A/L312A) LMX1B-FLAG. Levels are presented relative to the control signal (pcDNA 3.1 condition). Mean ± SEM (n = 3); one-way ANOVA followed by Tukey’s multiple comparison post-hoc test comparing scrambled sequence against NURR1 or pro-insulin: *** p<0.001 (LMX1B-FLAG) and *** p<0.001 (Y309A/L312A LMX1B-FLAG).

The release of the nuclear LC3 pool during starvation is regulated by SIRT1-dependent deacetylation, with LC3 exiting the nucleus in complex with the diabetes- and obesity-regulated nuclear factor (DOR) in preparation for its incorporation into the nascent isolation membrane (Huang et al., 2015, Mauvezin, Orpinell et al., 2010). GFP-TRAP pull-downs in HEK293T cells expressing acetylation-resistant (K49R/K51R) and acetylation mimic (K49Q/K51Q) GFP-LC3B (Huang et al., 2015) revealed that LMX1B (**Fig 5C**) and LMX1A (**Appendix Fig S3H**) associate strongly with K49R/K51R LC3B, but not with K49Q/K51Q LC3B, suggesting that LMX1A/LMX1B bind to the non-acetylated form of LC3B. The same was true for LIR-dependent LC3B binding to p62 (**Fig 5C; Appendix S3H**) and ULK1 (data not shown), indicating that the LC3B acetylation prevents LIR-type binding more generally. By contrast, non-LIR dependent binding to ATG3 (e.g. (Nakatogawa, 2013)) was retained—albeit noticeably reduced—in the K49Q/K51Q LC3B mutant (**Fig 5C**). Together, these data suggest that LMX1B (and LMX1A) binding to LC3B is regulated by LC3B acetylation/deacetylation, and that parallel LMX1B and LC3B nuclear shuttling itineraries exist, with the LMX1B association with LC3B being dependent on nutrient levels, sub-cellular localisation and LC3B acetylation status.

### LMX1B LIR mutation abolishes rotenone protection in human mDANs

Mutagenesis of the key tyrosine/leucine residues within the proposed LMX1B LIR motif (Y309A/L312A) markedly reduced binding to GFP-LC3B and GFP-GABARAP-L2 in GFP-TRAP pull-downs in HEK293T cells (**Fig 5D**), despite having negligible impact on its nuclear localisation (**Fig 5E**). Binding to GFP-LC3A, GFP-LC3C, GFP-GABARAP and GFP-GABARAP-L1 was apparently unchanged (**Fig 5D**), indicative of the presence of an alternative LIR(s) or due to stabilising interactions outside of the core LIR (Wirth et al., 2019). Consistent with this, lysosomal turnover of wild-type and LIR mutant LMX1B occurred at comparable rates in HEK293T cells (**Fig 5F**). Crucially, however, the protection afforded by LMX1B overexpression against rotenone toxicity in iPSC-derived human mDANs (see **Fig 3F**) was abolished when Y309A/L312A LIR mutant LMX1B was expressed (**Fig 5G**). This argues that LMX1B binding to LC3B (and potentially other ATG8 family members) is needed for its protective influence in human mDANs. Nuclear LMX1B-LC3B binding might enhance LMX1B-mediated protection via directed transcriptional modulation of autophagy pathway genes; however, knockdown/rescue experiments in HEK293T cells (**Fig 5H**) and in mDANs (**Fig 5I**) indicated similar levels of recovery of autophagy target gene expression by qRT-PCR (a possible exception being optineurin in mDANs). Consistent with this observation, Y309A/L312A LIR mutant LMX1B enhanced NURR1 and insulin FLAT (German, Wang et al., 1992) sequence-driven luciferase expression to the same extent as wild-type LMX1B (**Fig 5J**). Overall, these findings support a model in which cross-talk between LMX1B and the autophagy system is an important facet of cellular survival responses in cells mDANs, further emphasising the importance of understanding the roles and molecular control of LMX1B (and LMX1A) transcription factors in PD.

## DISCUSSION

A number of studies have highlighted the importance of autophagy quality control pathways in maintaining cell and tissue homeostasis and in the prevention of disease (see (Levine & Kroemer, 2008, Thorburn, 2018)). It is therefore hoped that strategies to maintain and/or boost autophagy levels in afflicted tissues will be beneficial in delaying and even preventing the onset of diseases, particularly those that are commonly associated with ageing (Rubinsztein, Marino et al., 2011). To realise its potential, the regulatory pathways that control tissues-specific autophagy responses must first be comprehensively understood. The existence of several classes of broad acting and/or tissue restricted transcription factors that act as autophagy regulators in different tissues, cell-types, and cellular contexts (Fullgrabe et al., 2016, Fullgrabe et al., 2014), provides a possible route to design strategies to boost autophagy responses in disease-susceptible cells and tissues (e.g. neurons).

The LIM homeodomain transcription factors LMX1A and LMX1B are essential for both the differentiation and the continued survival of mDANs in the substantia nigra region of the midbrain (Arenas, Denham et al., 2015, Doucet-Beaupre et al., 2016, Laguna et al., 2015, Nakatani et al., 2010). In addition to controlling expression of a number of downstream dopaminergic neuronal regulatory gene pathways, they sustain efficient autolysosomal quality control systems and regulate key nuclear-encoded mitochondrial genes in mDANs (Doucet-Beaupre et al., 2016, Laguna et al., 2015). Improving the stability and/or activity of LMX1A/LMX1B might therefore provide a route to boosting mDAN autophagy and mitochondrial function—and therefore mDAN resilience—in PD (Doucet-Beaupre et al., 2016). Here, we demonstrate that LMX1A and LMX1B are human autophagy transcription factors that each contribute to basal autophagy control and can both protect against PD-associated neuronal stress in vitro. Whilst a number of previous studies have highlighted the potential benefits of chemical/small molecule upregulation of autophagy in cell-lines and PD animal models (Harris & Rubinsztein, 2011, Malagelada, Jin et al., 2010, Renna, Jimenez-Sanchez et al., 2010, Sarkar, Davies et al., 2007, Tanji, Miki et al., 2015), our work is an important example of how autophagy upregulation through transcriptional control can have a positive impact in a disease-relevant context (human mDANs subjected to PD-associated stress). This parallels the protection demonstrated against α-synuclein toxicity following TFEB (or Beclin-1) overexpression in the rat midbrain in vivo (Decressac et al., 2013), with the added benefit of targeting regulatory pathways that are strongly associated with mDANs (Doucet-Beaupre et al., 2015). Intriguingly, we show that LMX1A and LMX1B factors interact with ATG8 family members via conserved LIRs, and in the case of LMX1B, this interaction appears to be needed for LMX1B’s protective roles in human mDANs in vitro.

Mutagenesis of the proposed LIR motif in LMX1B demonstrated that altering the two key residues—the aromatic phenylalanine at Θ_0_ and the aliphatic leucine at Г_3_—in the core LIR is sufficient to reduce binding to LC3B and GABARAPL2 (other ATG8 interactions were apparently unaffected). This suggests the presence of alternative LIRs in LMX1B, or that amino acids up- or downstream of the core LMX1B LIR may play substantial roles in ATG8 binding. In a recent study, Wirth et al. explored the significance of residues C-terminal to the core LIR in mediating interactions with GABARAPs (Wirth et al., 2019). The influence of amino acids C-terminal to the LMX1B LIR in mediating ATG8 interactions remains unclear, although we suggest that the glutamine at X_7_ may stabilise the interaction with LC3B, as is seen in the C-terminally-extended LIR in FYCO1 (Olsvik et al., 2015). Despite mutations in the core LIR not apparently affecting binding to several ATG8 family members, the mDAN protection afforded by overexpression of wild-type LMX1B was not observed using this construct. This suggests that the LMX1B interactions with LC3B or GABARAPL2 may be of particular importance. Surprisingly, we were unable to detect obvious differences in autophagy gene expression in knockdown/rescue experiments using wild-type and LIR mutant LMX1B in HEK293T cells and in mDANs. It is therefore unlikely that altered LMX1B-mediated autophagy gene expression due to differential binding to LC3B and/or GABARAPL2 explains the protective influence of LMX1B against stress. Since LMX1B autophagy turnover rates were not affected by core LIR mutations, it is possible that the location of the LMX1B-ATG8 interaction is important. LC3B is known to cycle between nucleus and cytosol in response to nutrient stress, with deacetylation at key lysines adjacent to the LIR-docking site and DOR-associated nuclear import needed for establishing LC3B lipidation competency (Huang et al., 2015). Interestingly, the LMX1B-LC3B interaction took place exclusively in the nucleus in fed cells, but was dramatically increased in the cytosol during starvation, when clear co-localisation between LMX1B and LC3B in cytosolic puncta could be observed. This suggests that, like LC3B, LMX1B undergoes nutrient-dependent shuttling between nucleus and cytoplasm. One obvious reason for this is to enable autophagy-mediated degradation of LMX1B during prolonged stress; however, a role for LMX1B in coordinating LC3B nuclear export cannot be ruled out.

In conclusion, we have described a LIR-dependent interaction between the autophagy transcription factors LMX1A and LMX1B and the core autophagy machinery. We propose that this represents a novel autophagy regulatory loop where transcription factors that control autophagy gene expression are themselves degraded by autophagy to potentially restrict the duration and/or strength of the autophagy response. This finding has implications for our understanding of the control of autophagy during chronic stress in disease-susceptible tissues.

## MATERIALS AND METHODS

### Resources table

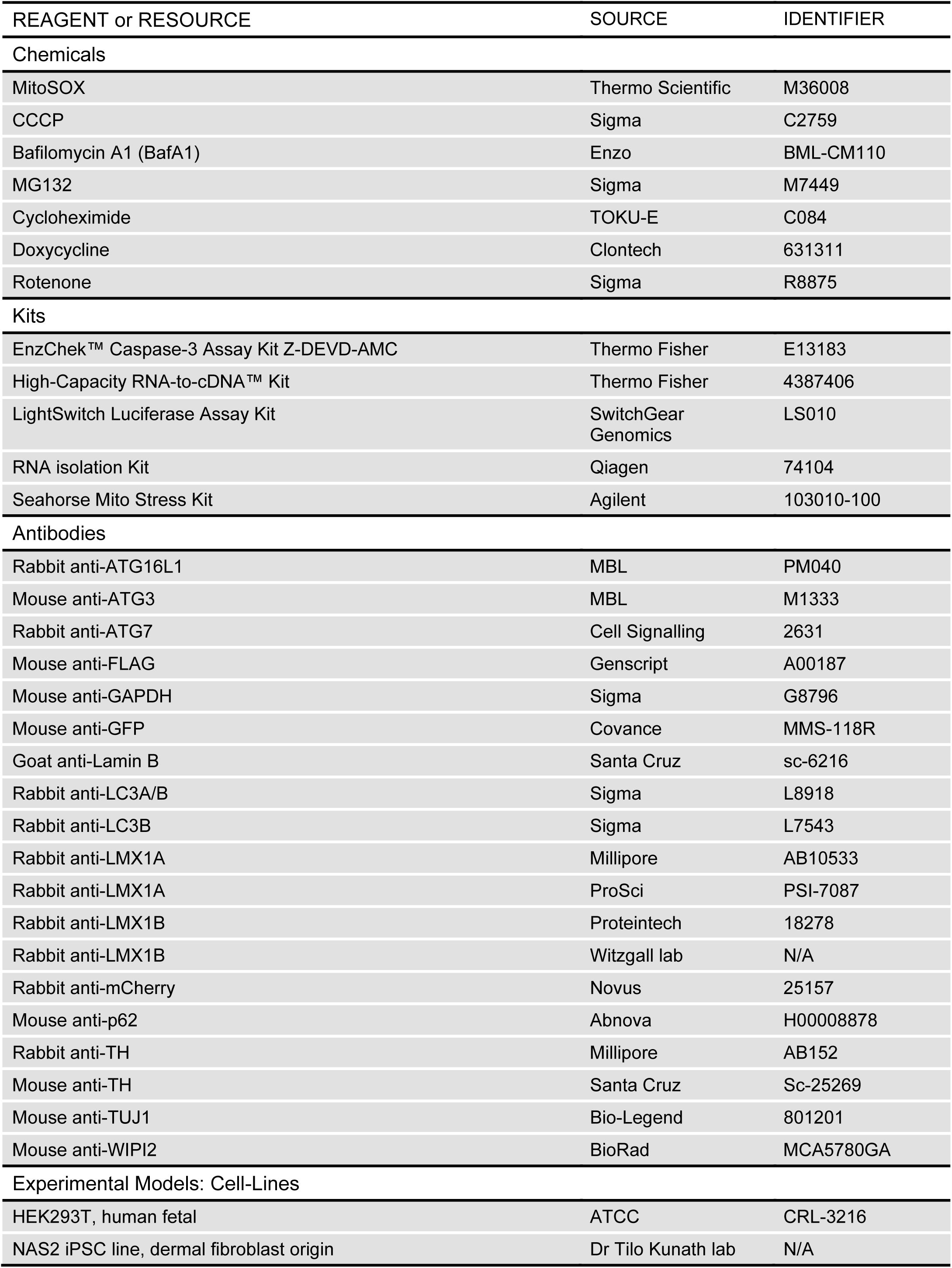
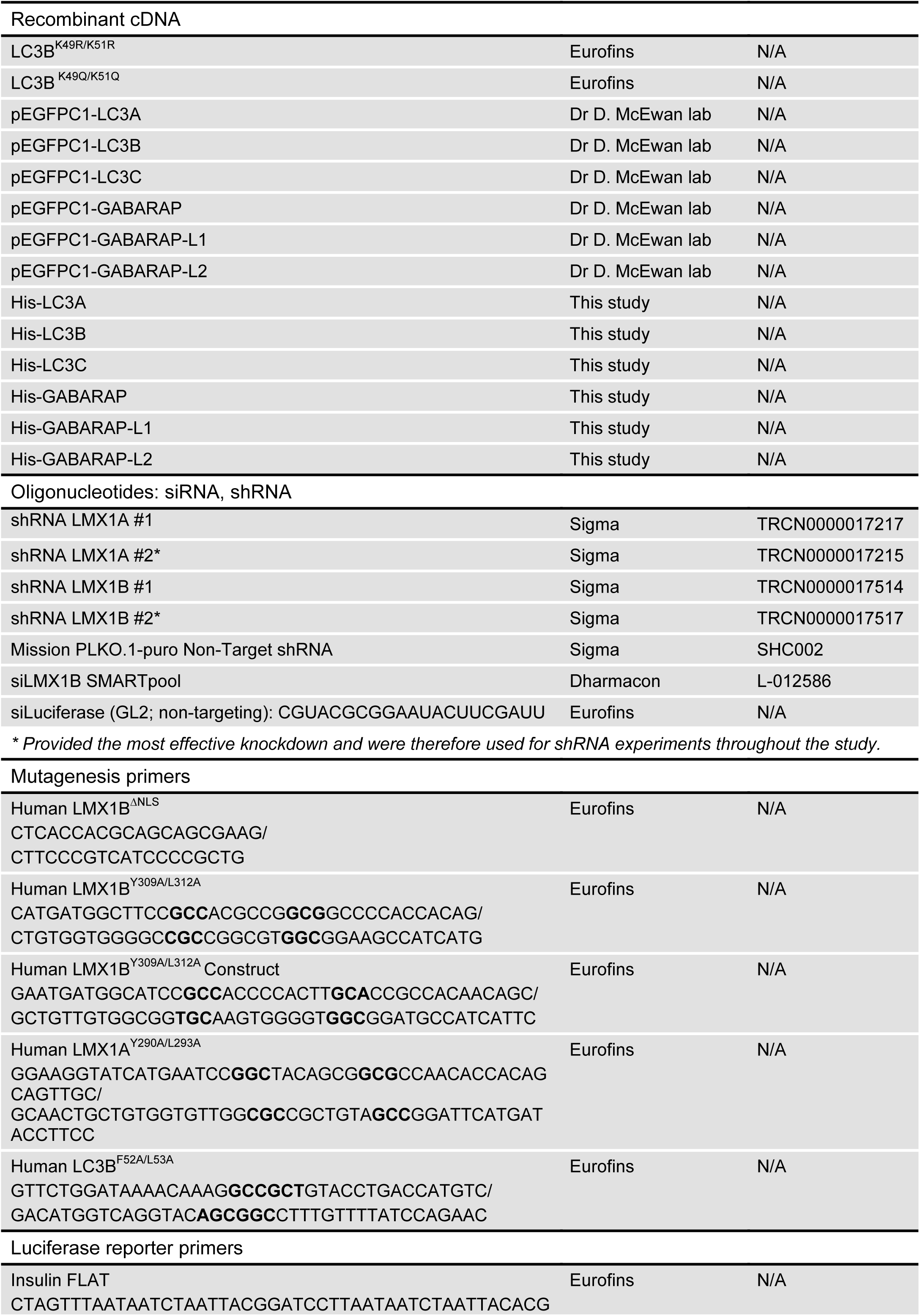
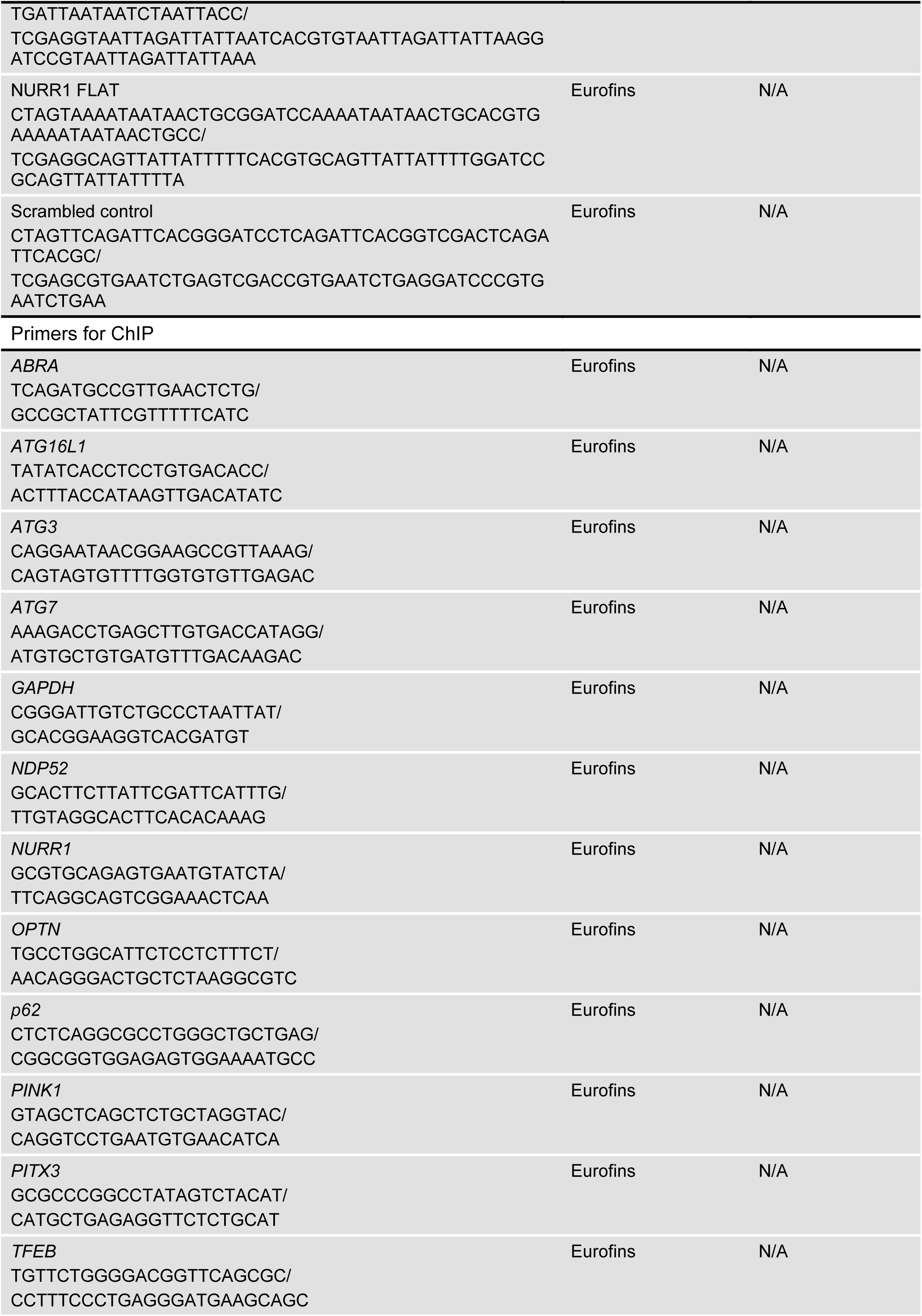
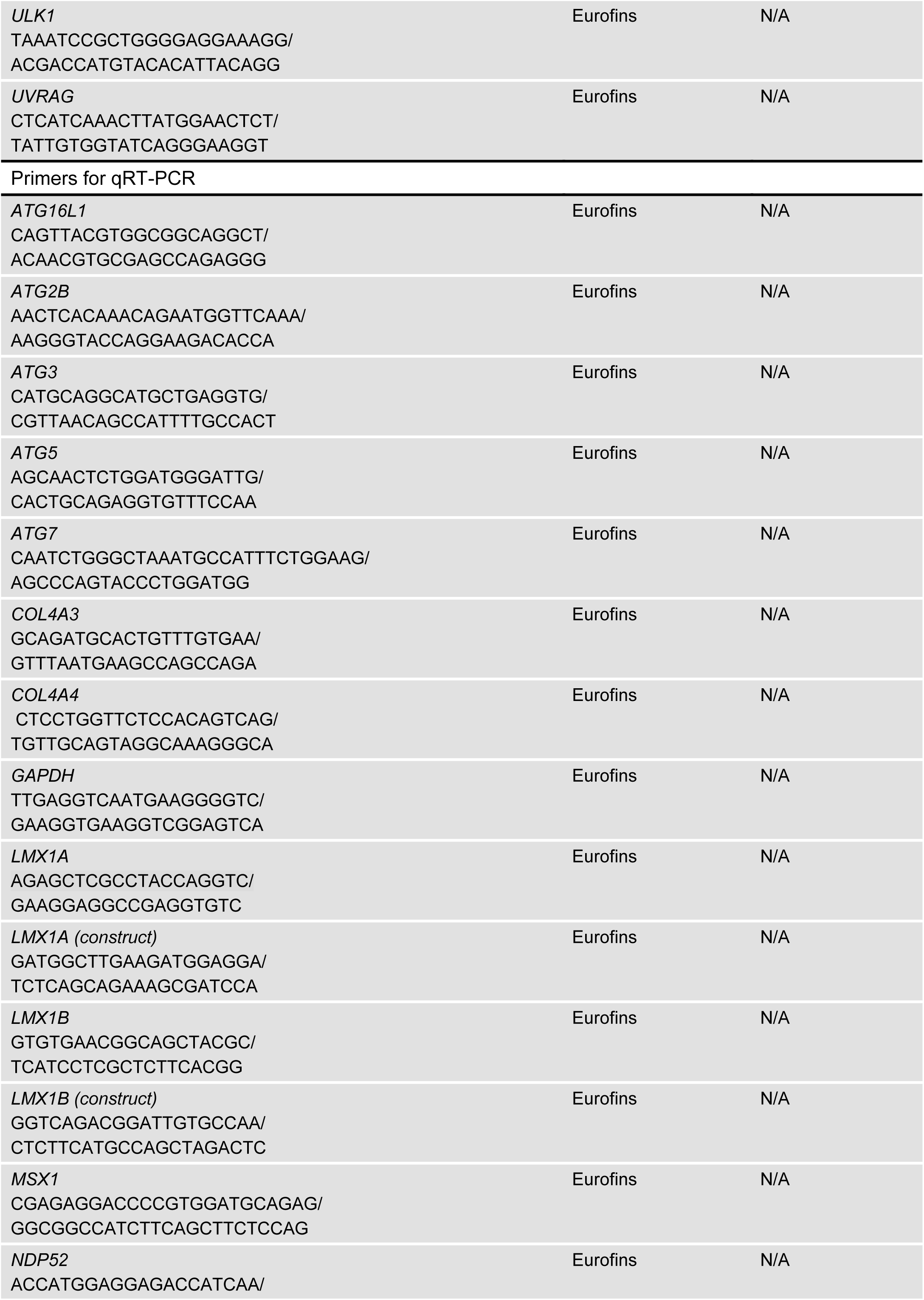
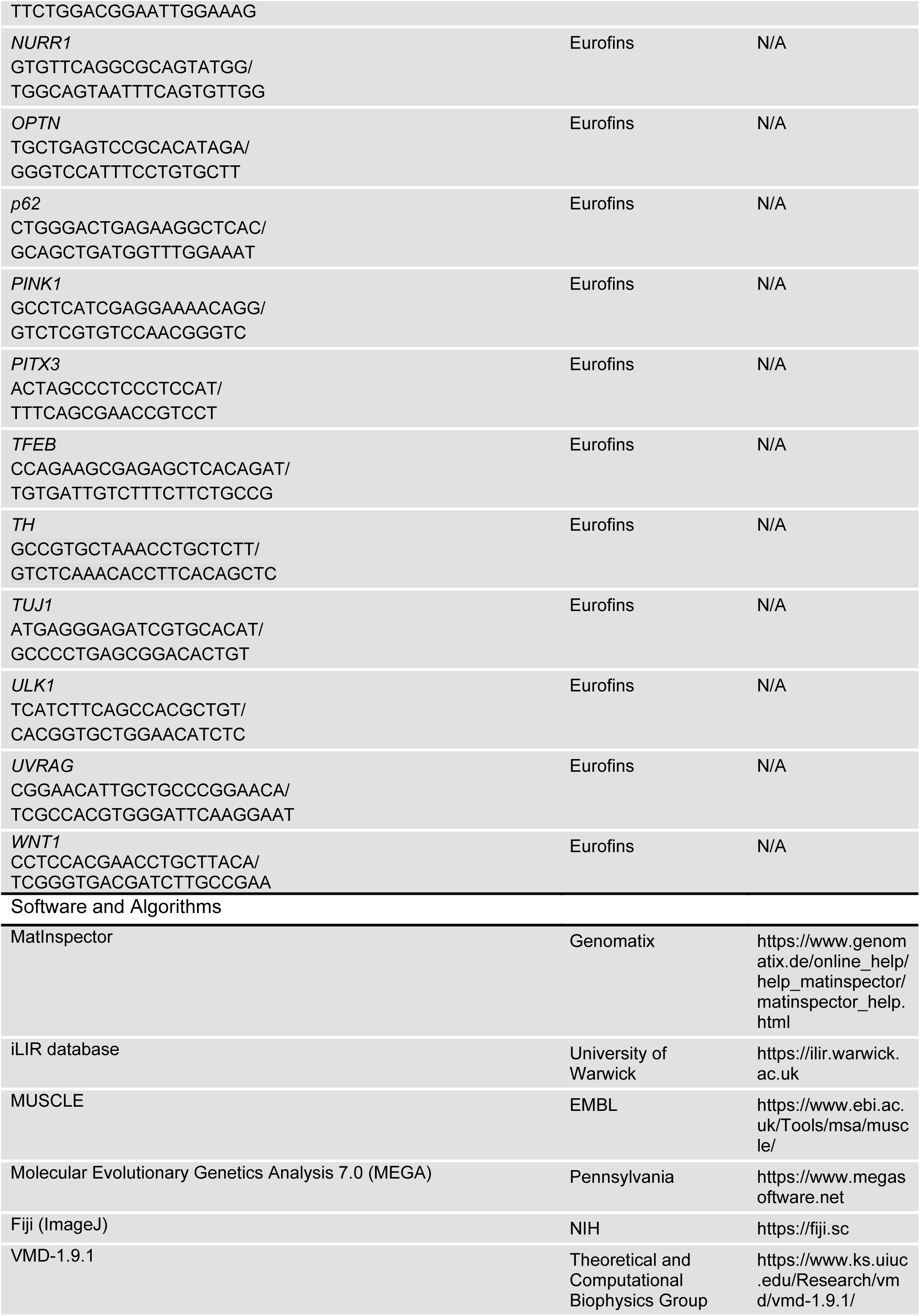
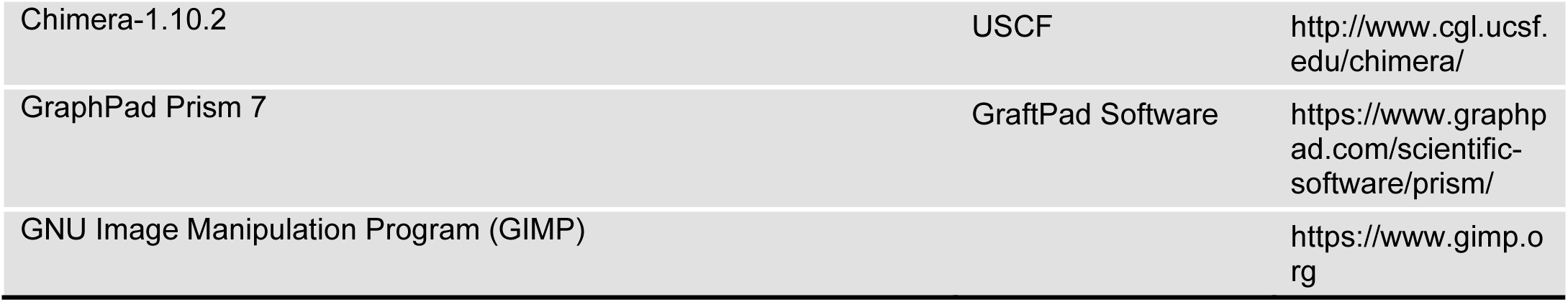

### Antibodies

The following primary antibodies were used (key: IB, immunoblot concentration; IF, immunofluorescence concentration; IP, used for immunoprecipitation): anti-ATG16L1 (MBL, PM040: IB, 1:1000; IF, 1:400); anti-ATG3 (MBL, M1333: IB, 1:1000); anti-ATG7 (Cell Signalling, 2631: IB, 1:1000); anti-FLAG (Genscript, A00187: IB, 1:1500; IF, 1:300); anti-GAPDH (Sigma, G8796: IB, 1:2000); anti-GFP (Covance, MMS-118R: IB, 1:2000); anti-Lamin B (Santa Cruz, sc-6216: IB, 1:500); anti-LC3A/B (Sigma, L8918: IB, 1:1000); anti-LC3B (Sigma, L7543: IF, 1:400; IP); anti-LMX1A (Millipore, AB10533: IB, 1:1000; IF, 1:1000; IP); anti-LMX1A (ProSci,PSI-7087: IP); anti-LMX1B (Proteintech, 18278: IB, 1:1000; IF, 1:250; IP); anti-LMX1B (Witzgall lab: IB, 1:2000; IF, 1:350); anti-p62 (Abnova, H0008878: IB, 1:1000; IF, 1:400); anti-mCherry (Novus, 25157; IB, 1:1000); anti-TH (Millipore, AB152: IB, 1:1000; IF, 1:300); anti-TH (Santa Cruz, sc-25269: IF, 1:100); anti-TUJ1 (Bio-Legend, 801201: IB, 1:1000; IF, 1:1000); anti-WIPI2 (BioRad, MCA5780GA: IF, 1:400). Secondary antibodies for immunoblotting were: anti-mouse HRP (Stratech, G32-62DC-SGC: 1:10000), anti-rabbit HRP (Stratech, G33-62G-SGC: 1:10000) and anti-goat HRP (Stratech, G34-62DC-SGC). Secondary antibodies for immunofluorescence were: Alexa Fluor 488 (Invitrogen, A-11029, A-11034: 1:400), Alexa Fluor 568 (Invitrogen, A-11031, A11036: 1:400) and Alexa Fluor 647 (Invitrogen, A-21236, A-21244: 1:400).

### Culturing immortalized cells

HEK293T human embryonic kidney cells were maintained in high-glucose DMEM medium supplemented with 10% FBS (Sigma) at 37°C in 5% CO_2_. Transient transfections were performed with Lipofectamine 2000 Reagent (Thermo Scientific). For luciferase experiments, cells were cultured in DMEM phenol-red free medium supplemented with 10% FBS (Thermo Fisher). For starvation experiments, the following media was used: 140mM NaCl, 1mM CaCl_2_, 1mM MgCl_2_, 5mM glucose and 20mM HEPES supplemented with 1% BSA (Sigma) (Axe, Walker et al., 2008).

### iPSC culture

We used the wild type α-synuclein 2 (NAS2) iPSC line derived from human dermal fibroblasts by retroviral reprogramming at passages 40-80 (kindly provided by Dr. Tilo Kunath, Center of Regenerative Medicine, University of Edinburgh, United Kingdom) (Devine et al., 2011). NAS2 were maintained in Essential 8 TM Medium (E8) supplemented with Essential 8 Supplement (Thermo Scientific) and RevitaCell (1/100, Thermo Scientific) in plates previously coated with 5μg/mL Vitronectin (in PBS; Thermo Scientific) at 37°C in 5% CO_2_ as fully described in (Stathakos et al., 2019).

### Midbrain dopaminergic neuronal differentiation and maintenance

We used an improved monolayer protocol for mDAN differentiation based on previously published protocols (Nistor, May et al., 2015, Torper, Pfisterer et al., 2013), and described in (Stathakos et al., 2019). mDAN differentiation was achieved by the addition of SMAD inhibitors and the pattering factors, WNT and SHH (Arenas et al., 2015). iPSCs in small colonies (commonly three days after initial plating) were grown in N2B27 neural differentiation media for 9 days, comprising 50% neurobasal media (Thermo Scientific) and 50% DMEM/F-12 with Glutamax (Thermo Scientific), supplemented with: N2 (1/200, Thermo Scientific); B27 (1/100, Thermo Scientific); 1mM Glutamax (Thermo Scientific); 5mg/mL insulin (Sigma); nonessential amino acids (1/100 Thermo Scientific); Penicillin/Streptomycin (1/100, Sigma); 75mM 2-mercaptoethanol (Thermo Scientific); neural induction factors, LDN (100nM; Sigma) and SB431542 (10μM; Tocris); and patterning factors, SHH (200ng/mL; R&D Systems) and WNT homolog, Chir (0.8μM; AxonMedchem). Cells were passaged using StemPro Accutase (Thermo Scientific), and typically plated at 1:2-1:3 dilution ratio in neural differentiation media supplemented with RevitaCell on plates coated with poly-L-ornithine (Sigma) and laminin (1/1000 in PBS, Sigma). At day 9 of neuralization, induction and patterning factors were removed. After the next passage (usually at day 11), the N2B27 media was supplemented with the neurotrophic factors: 20ng/mL brain-derived neurotrophic factor (BDNF; Peprotech), 20ng/mL glial cell-line-derived neurotrophic factor (GDNF; Peprotech), and 0.2mM ascorbic acid (AA; Sigma). For terminal differentiation/maturation, cells were passaged on poly-L-ornithine/laminin coated coverslips or plates, and cultured in complete N2B27 supplemented with db-cAMP (0.5mM; Sigma) and N-N-(3,5-difluorophenacetyl)-1-alanyl-S-phenylglycine t-butyl ester (DAPT; 5μM; Tocris) at 37°C in 5% CO_2_ for 7-14 days depending on the experiment.

### Plasmids and transfection

To overexpress LMX1A/B, either the Tet-On 3G inducible system (Clontech), LMX1B-FLAG in pcDNA3.1 (Genscript), or pLVX-puro plasmids were used. The Tet-on 3G system is based on two plasmids: (i) the CMV-Tet plasmid expressing the Tet-On transactivator protein, and (ii) TRE3G promoter controlling the expression of a gene of interest, in this case, LMX1A (NM_001174069.1) and LMX1B (NM_001174147.1)–empty plasmid was used as a control. cDNAs were synthetized codon optimized for H. sapiens (Eurofins Genomics). Doxycycline was added for 48-72h. For inducible expression of LMX1A/B in iPSC-derived neurons, the human synapsin promoter (hsyn) in pRRL plasmid (kindly provided by Prof. James Uney, University of Bristol) was used to drive expression of the Tet-On activator protein. The pRRL plasmid was also used as a backbone to generate shRNA reporter constructs: hsyn-GFP-U6-shRNA. Human ATG8 family members in pEGFPC1 were provided by Dr David McEwan (Dundee, UK). Plasmid sequences are available on request.

### Viruses, transduction and stable cell lines

Lentivirus were produced in HEK293T cells by transient transfection using PEI reagent (Sigma). 27µg of the plasmid of interest was transfected together with 20.4µg of the packing plasmid pAX2 and 6.8µg of the envelope plasmid pMGD2. Viruses were harvested 48h after transfection. Media was collected and centrifuged 1,500*g* for 5min and filtered with a 0.45µm filter to remove cells and debris. Viruses were concentrated using Lenti-X Concentrator (Clontech). One volume of Lenti-X Concentrator was combined with three volumes of clarified supernatant. The mixture was incubated 1h at 4°C, then centrifuged at 1,500*g* for 45min at 4°C and pellet was resuspended in N2B27 media or DMEM media. For viral transduction, neural progenitors were infected for 3 days with 3µl viruses/mL of media and then media was removed and replace with complete N2B27 for another 2-4 days. The efficacy of transfection was checked using fluorescence microscopy.

HEK239 cells overexpressing LMX1B-FLAG, and HEK293 cells overexpressing mCherry-LMX1B in a Tet-On 3G system were generated as follows: cells were plated in 6cm dishes and transduced with the corresponding lentiviruses in the presence of 10µg/mL polybrene (Sigma) to increase transduction efficiency. After two days, cells were fed with media supplemented with 1µg/mL puromycin—for non-inducible expression—together with 800µg/mL G418—for the Tet-On 3G system. These concentrations were selected using a titration kill curve in HEK293 (data not shown). Cells were fed every 3 days. For Tet-On 3G stable cell lines, prior to the experiment, 500ng/mL doxycycline was added to the media for 48h to induce the expression of the gene of interest.

### Short interfering RNAs (siRNA) transfection

In HEK293T cells, siRNA transfection was carried out through reverse transfection protocol with lipofectamine. 3µl of 20µM siRNAs were mixed with 2µl lipofectamine in Opti-MEM reduced-serum media (Thermo Scientific) and combine with 300×10^3^ cells per well of a 6-well plate. Cells were plated overnight in Opti-MEM media. The following day, cells were fed with DMEM + FBS media. A second forward transfection step was then carried out with the reagents described above. Samples were collected 48h after the second transfection. For rescue experiments, the LMX1B cDNA was synthesised codon optimized for *H. sapiens*, so that the siRNA would not recognize the targeting sequence (Eurofins Genomics), and this was cloned into the pLVX plasmid (empty pLVX plasmid was used as a control).

### Luciferase assay

For luciferase assay, the LightSwitch™ Vector (SwitchGear Genomics) was digested with NheI and XhoI (Biolabs) for the cloning of the promoters. 3x tandem repeats of the putative FLAT elements for pro-insulin (German et al., 1992) and NURR1–a scramble sequence was used as a control (Pajares, Rojo et al., 2018)–connected by linker regions with a BamHI site. Primers were annealed and ligated with the digested promoter with T4 ligase (Biolabs). Cells were seeded on CELLSTAR® 96-well white plate (Greiner Bio-one) in DMEM phenol-red free medium at a density of 60-70%. The following day, cells were transfected with LightSwitch™ Vector with a FLAT-containing promoter—or a scrambled sequence as control—together with pcDNA3.1 LMX1B-FLAG (or LMX1B^Y309A/L312A^-FLAG)—or pcDNA 3.1 as control. Plasmids were transfected in a ratio of 0.1µg/each plasmid: 0.45µl transfection reagent— lipofectamine (HEK293T). After 24h, plates were frozen at −80°C to increase luciferase signal. To measure luciferase, the LightSwitch Luciferase Assay Reagent assay (SwitchGear Genomics) was used according to manufacturer’s instructions. Luciferase levels were measured in a Fusion Universal microplate reader (PerkinElmer) with a Photomultiplier tubes (PMT) voltage of 1100 and each well was read for 2secs. Relative luciferase levels were normalized to luciferase control signal (pcDNA 3.1 condition).

### GFP-trap immunoprecipitation

HEK293 cells were plated on 10cm dishes co-transfected with the corresponding GFP-tagged constructs and LMX1A or LMX1B-FLAG (or the LIR mutated versions). DNA was transfected using 1mg/mL PEI reagent in a ratio of 1:6 in Opti-MEM. Cells were washed twice with PBS, then lysed in 500µl of GFP-trap lysis buffer containing: 50 mM Tris–HCl, 0.5% NP40, 1mM PMSF, 200µM Na_3_VO_4_ and 1/50 protease inhibitor tablets (Roche). The samples were incubated on ice for 10min, then lysates were slowly forced through a 20G needle to help break the nuclei. Soluble fractions were obtained by centrifugation at 13,000*g* for 10min at 4°C. 5% of the sample was kept as total lysate sample. The remainder was incubated with 20µl GFP-trap beads (Chromotek), previously washed with GFP-trap wash buffer containing 50 mM Tris-HCl, 0.25% NP40, 1mM PMSF, 200µM Na_3_VO_4_ and 1/50 protease inhibitor tablets, for 2h at 4°C. Then, beads were washed three times with GFP-trap wash buffer and a fourth time with GFP-trap wash buffer 2: 50 mM Tris–HCl, 1mM PMSF, 200µM Na_3_VO_4_ and 1/50 protease inhibitor tablets. Beads were then resuspended in 30µl 2x loading buffer.

### Recombinant protein purification and in vitro binding assays

Human ATG8 sequences were cloned into His-tag plasmid (ptrHisC), and these were transformed into E. coli BL21 (DE3) bacteria (Biolabs). To induce expression, 0.5mM IPTG (Sigma) was added for 3h. The culture was centrifuged at 1,500*g* for 10 min 4°C, and the pellet resuspended in 13mL homogenization buffer comprising: 25mM Tris pH 7.5, 1% Triton X-100, 250mM NaCl, 20mM imidazole (Sigma). The sample was sonicated on ice using a sequence of 10sec on / 20sec off for 5 min. The soluble fraction was harvested after centrifugation at 3,000*g* for 30min at 4°C. and incubated with 1ml nickel-chelating Resin (Probond) previously washed twice with homogenization buffer (1ml resin/10 ml buffer) for 1h at 4°C. Poly-Prep® Chromatography Columns (Bio-Rad) were used to pack the resin before elution. The column was extensively washed with homogenization buffer without imidazole, and His-ATGs were eluted with 500µl elution buffer (20mM Tris pH 7.5, 1M NaCl, 100mM EDTA, 200mM imidazole).

Recombinant ATG8 proteins were then coupled to cyanogen bromide-activated sepharose 4 Fast Flow (CNBr-sepharose, GE Healthcare) as described (Kavran & Leahy, 2014). First, the recombinant proteins were dialyzed using Slide-A-Lyzer™ cassettes (Thermo Fisher) into cold coupling buffer containing 100mM NaHCO_3_ and 500mM NaCl. To activate the resin, 0.25g of resin was incubated with 5 volumes of 1mM HCl for 2h at 4°C producing 1ml of hydrated resin. Resin was washed with 1mM HCl, then 2mg of recombinant protein (measured using nanodrop A280) was coupled to 1ml of hydated resin overnight at 4°C. As a negative control, blank resin without incubation with recombinant protein was prepared to test for unspecific binding. The reaction was quenched by incubation with quenching buffer containing 100mM Tris-HCl pH 7.5 for 3h 4°C. Uncoupled protein was removed by washing with high pH / low pH wash buffers comprising: 100mM Tris-HCl pH8, 500mM NaCl / 100mM NaOAc, 500mM NaCl. For in vitro binding assays, HEK293T cells grown to sub-confluency on 10cm dishes were washed with ice-cold PBS and lysed with 500µl of RIPA buffer supplemented with protease inhibitor. The samples were incubated on ice for 10min and then lysates were diluted 1:2 with CNBr IP wash buffer (50 mM Tris–HCl pH 7.5, 150mM NaCl, 1mM EDTA, 1/50 protease inhibitor tablets [Roche]). The homogenates were incubated on ice for 15min and were cleared by centrifugation at 12,000*g* for 15min at 4°C. Prior to the incubation, resin was washed in CNBr IP wash buffer. For the binding, 250µl of the cleared lysate diluted with 140µl of CNBr IP wash buffer was incubated with 30µl of the coupled resin overnight 4°C. Then, beads were washed three times with CNBr IP wash buffer, then resuspended in 40µl 2x loading buffer.

### Cell fractionation

HEK293 cells were plated on 10cm dishes and transfected with the LMX1B-FLAG or LMX1A with/without GFP-ATG8s cDNAs using 1mg/mL PEI reagent at a ratio of 1:6 in Opti-MEM for 24h. Cells were washed twice in 5mL of ice-cold PBS and harvested by centrifugation at 200*g* for 5min 4°C. The cell pellet was then lysed with 400µl of Buffer A [20mM HEPES pH 7 (Sigma), 0.15mM EDTA, 0.015mM EGTA (Sigma), 10mM KCl (Sigma) and 1% NP-40 (Sigma) supplemented with one tablet of protease inhibitor per 10mL of buffer, as described in (Garcia-Yague, Rada et al., 2013)] and incubated on ice for 30min. The homogenate was centrifuged at 1,000*g* for 5min 4°C, after which the supernatant was collected, and the nuclear pellet washed in 500µl Buffer B (10mM HEPES pH 8, 25% glycerol, 0.1M NaCl, 0.1mM EDTA supplemented with one tablet of protease inhibitor per 10mL of buffer). After centrifugation as above, the nuclear pellet was incubated with DNaseI (Thermo Fisher) in 100µl of Buffer A for 20min and then 4x loading buffer was added to the sample. If the nuclear fraction was required for GFP-trap immunoprecipitation, after washing with Buffer B, it was resuspended in 300µl of Buffer C (10mM HEPES pH 8, 25% glycerol, 0.4M NaCl and 0.1mM EDTA supplemented with one tablet of protease inhibitor per 10mL of buffer) for 30min at 4°C. Samples were centrifuged at 4,500*g* 20min 4°C with soluble nuclear proteins located in the supernatant. 120µl of this fraction was kept as total nuclear fraction and the rest of the fraction was diluted up to 600µl with GFP-trap wash buffer for the incubation with GFP-trap beads.

### Protein turnover experiments

Stable HEK293 LMX1B-FLAG cells were plated on 6-well plates at 70-90% confluency. Cells were treated for 16h with cycloheximide (50µg/mL) in the absence or presence of BafA1 (20nM) or MG132 (10µM). For starvation experiments, cells were treated for 6h with cycloheximide (50µg/mL) in starvation media in the absence or presence of BafA1 (20nM). Lysates were collected for immunoblotting.

### Immunoblotting

Cells grown on 6-well plates were initially washed with ice-cold PBS, then lysed with 100-200µl/well of ice-cold radioimmunoprecipitation assay (RIPA) buffer consisting of 50mM Tris HCl (pH7.4), 1% Triton-X-100 (Sigma), 0.5% sodium doexycholate (Sigma), 150mM NaCl (Sigma), 0.1% sodium dodecyl sulphate (SDS; Sigma) supplemented with one tablet of protease inhibitor per 10mL of RIPA buffer. The homogenates were incubated on ice for 15min, then cleared by centrifugation at 12,000*g* for 15min at 4°C. Supernatants were collected as soluble fractions. Sample protein concentration was determined by BCA protein assay (Pierce) according to manufacturer’s protocol. Proteins were transferred to nitrocellulose membranes (Biolabs). Membranes were then incubated with primary antibody diluted in 2.5% milk or 2.5% BSA in Triton X-100-TBS buffer (T-TBS) for 2h or overnight. Primary and secondary antibodies used are listed above. Membranes were then washed three times prior to incubation with ECL Chemiluminiscence reagents (Geneflow), and band intensities were detected in films (GE Healtchare) using a film developer.

### Immunofluorescence

Cells were seeded on coverslips (pre-coated with poly-L-ornithine and laminin for neurons). iPSC-derived mDANs were allowed to mature for 7-11 days prior to fixation. Cells were washed twice with PBS and incubated with 4% formaldehyde (Thermo Scientific) for 20min or −20°C methanol for 5min. Formaldehyde-fixed cells were incubated with blocking solution containing 5% BSA and 0.3% Triton X-100 (Sigma) in PBS for at least half an hour at room temperature. Cells were then incubated 2h/overnight with primary antibody (listed above) prepared in 2.5% BSA (including 0.15% Triton X-100 for neurons). Cells were washed three times with PBS and incubated with the secondary antibodies (listed above) and counterstained with DAPI prepared in 2.5% BSA (and 0.15% Triton X-100 for neurons) for 1h. Cells were then washed again with PBS, mounted in Mowiol, and fluorescence images were captured using a Leica DMI6000 SP5-II confocal microscope. Image analysis was performed using Fiji (Schindelin, Arganda-Carreras et al., 2012) (National Institutes of Health, Bethesda, USA).

### Chromatin immunoprecipitation (ChIP)

HEK293T cells or iPSC-derived mDANs were plated on 10cm dishes at 80-90% confluency. ChIP assays were conducted based on previously published protocols (Pescador, Cuevas et al., 2005). Firstly, cells were fixed with 625µl 16% formaldehyde added to the media (Thermo Fisher) on ice for 12min and the crosslinking reaction was stopped with 125mM glycine (Sigma). Cells were then washed twice with ice-cold PBS supplemented with protease inhibitors and harvested by centrifugation at 200*g* for 5min at 4°C. The pellet was lysed with 200µl with ChIP lysis buffer (1% SDS, 10mM EDTA and 50mM Tris pH 8.1 supplemented with protease inhibitors) and incubated on ice for 10min. Samples from different dishes were pooled for sonication on ice using a probe sonicator with a sequence of 15sec on / 15sec off for 8min/plate to obtain an adequate fragment size of DNA (800-200bp). The homogenates were cleared by centrifugation at 12,000*g* for 10min at 4°C, and the supernatants collected as soluble fractions. Samples were then diluted in 10 volumes of ChIP dilution buffer comprising 0.01% SDS, 1.1% Triton X-100, 1.2 mM EDTA, 16.7mM Tris pH 8.1 and 167mM NaCl supplemented with protease inhibitors. The lysates were then precleared for 1h at 4°C using 10µl protein A agarose 50% slurry/plate (Millipore), and collected by centrifugation at 300*g* for 3min at 4°C (100µl of sample/plate was kept as input chromatin). The remaining lysate (∼2mL/plate) was used per ChIP experiment (approximately 1×10 cm dish per IP). Each sample was incubated overnight at 4°C with 4µg of anti-LMX1B (Proteintech), anti-LMX1A (PriSci) or 4µg of rabbit IgG (Cell Signalling). Immunocomplexes were recovered by incubation with 60µL pre-washed protein A/sample for 1h at 4°C. Prior to the elution of DNA, samples were incubated sequentially in the following buffers to improve the removal of non-specific chromatin interactions: (1) ChIP low salt buffer containing 0.1% SDS, 1% Triton X-100, 2mM EDTA, 20 mM Tris pH 8.1 and 150mM NaCl; (2) ChIP high salt buffer containing 0.1% SDS, 1% Triton X-100, 2mM EDTA, 20 mM Tris pH 8.1 and 500mM NaCl; (3) ChIP lithium buffer containing 1% Igepal (Sigma), 1mM EDTA, 10mM Tris pH 8.1, 250mM LiCl and 1% sodium deoxycholate; and (4) twice in TE wash buffer containing 10mM Tris pH 8 and 1mM EDTA. Finally, eluted samples were obtained by incubation with 250µl of ChIP elution buffer containing 0.1M NaHCO_3_ and 1% SDS for 15min at room temperature (this process was repeated twice so the total eluate was 500µl). Input chromatin samples were also diluted in ChIP elution buffer (500µl final volume). To reverse the cross-linking, all samples (including input chromatin) were incubated with 20µl 5M NaCl at 65°C overnight. Then, 10µl 0.5M EDTA, 20µl 1M Tris pH 6.5 and 2µl proteinase K (Thermo Fisher) was added to each sample and samples were incubated for 1h at 45°C. For the purification of DNA, the phenol-chloroform purification was used. Finally, the DNA pellet was resuspended in a final volume of 30µl and diluted 1:3 for qRT-PCR analysis with specific primers designed with the information obtained by bioinformatics analysis using the UCSC genome browser (California, USA). After the corresponding treatment/transduction, cells were washed with PBS and then cells were lysed in 350µl RLT buffer (Qiagen). Total RNA was extracted through columns using RNeasy kit (Qiagen) following manufacturer’s instructions and genomic DNA was digested using DNaseI (Qiagen). RNA samples were reverse transcribed using High-Capacity RNA-to-cDNA™ Kit (Thermo Scientific), according to manufacture’s protocol. The cDNA samples were amplified using SYBR Green (Life Technologies). The reaction was carried out using StepOnePlus System (Applied Biosystems) and the following conditions were selected: after an initial denaturation at 95°C for 10 min, 40 cycles with 95°C for 15s (denaturation), 60°C for 30s (annealing) and 60°C for 30s (elongation). For the analysis, mRNA levels were estimated using the ΔΔCt method and were normalised to GAPDH (Livak & Schmittgen, 2001).

### Analysis of mRNA levels by quantitative real-time polymerase chain reaction (qRT-PCR)

HEK293 cells and iPSC-derived mDANs were plated on 6-well or 12-well plates, respectively. Cells were allowed to mature for 3 days before transducting with the corresponding viruses for 3 days. Then, viruses were removed and replaced with complete N2B27 for another 2-4 days. After the corresponding treatment/transduction, cells were washed with PBS and then cells were lysed in 350µl RLT buffer (Qiagen). Total RNA was extracted through columns using RNeasy kit (Qiagen) following manufacturer’s instructions and genomic DNA was digested using DNaseI (Qiagen). RNA samples were reverse transcribed using High-Capacity RNA-to-cDNA™ Kit (Thermo Scientific), according to manufacture’s protocol. The cDNA samples were amplified using SYBR Green (Life Technologies). The reaction was carried out using StepOnePlus System (Applied Biosystems) and the following conditions were selected: after an initial denaturation at 95°C for 10 min, 40 cycles with 95°C for 15s (denaturation), 60°C for 30s (annealing) and 60°C for 30s (elongation). For the analysis, mRNA levels were estimated using the ΔΔCt method (Livak & Schmittgen, 2001) normalising data to GAPDH levels.

### IncuCyte cell imaging

iPSC-derived mDANs were seeded to mature in 24-well imaging plates (Thermo Fisher), and were transduced with hsyn-GFP/mcherry-U6-shRNAs viruses. Cells were imaged using the IncuCyte® S3 Live-Cell Analysis System (Essen Bioscience) at 20x magnification, with images obtained every 1h by phase contrast, GFP and mCherry fluorescence over a period of 3 days. Analysis was carried out by Dr Stephen Cross (Wolfson facility, University of Bristol). Analysis of neurite length and fluorescence intensity was performed in Fiji, using the Modular Image Analysis plugin (Cross, 2017). First, regions of each phase contrast image corresponding to cell bodies were removed using a mask image. Each mask was created by applying a variance filter to the phase contrast channel image, which enhanced large objects with high contrast, such as cell bodies. The filtered image was subsequently binarised using the Huang method (Xiao, Cao et al., 2011) (set to 80% its absolute value), holes in the mask filled and a median filter applied to smooth the object borders. The mask was applied to the original phase contrast image such that the masked regions had intensity close to the phase contrast image to minimise false object detection at the mask boundaries. Neurites were identified in the unmasked regions using the RidgeDetection plugin for Fiji with a minimum neurite length of 15px. The length of each neurite was measured, along with the green and red fluorescence channel intensities coincident with each neurite.

### Seahorse bioenergetics

Day 30-50 iPSC-derived mDANs were plated according to manufacturer’s instructions on 8-well Seahorse XFp plates (Agilent) previously coated with Poly-L-ornithine and laminin. Cells were allowed to mature for 3 days before being infected with hsyn-GFP-U6-shRNAs viruses for 3 days. Viruses were then removed, and media replaced with complete N2B27 for another 2-4 days. The day of the assay, culture media was replaced with Seahorse XF base medium (Agilent) supplemented with 1mM sodium pyruvate, 2mM glutamine, and 10mM glucose (pH 7.4) for 1h at 37°C. The Mito Stress Test Kit (Agilent) was prepared according to the manufacturer’s instructions: oligomycin (1µM); carbonilcyanide ptriflouromethoxyphenylhydrazone (FCCP; 1µM); rotenone/antimycin A (0.5µM). After analysis, cells were lysed in 20µl of RIPA buffer and protein levels were quantified by Nanodrop (A280) to normalise the data.

### Caspase assay

iPSC-derived mDANs were matured for 3 days in 96-well plates, then transduced with viruses for a further 3 days. Media was replaced with complete N2B27 with doxycycline (500ng/mL) for another 2-3days. Rotenone (15µM) or DMSO were added for 24h, and cells washed with PBS before plates were frozen at −80°C. Caspase levels were measured using the EnzChek™ Caspase-3 Assay Kit Z-DEVD-AMC substrate (Thermo Fisher) according to manufacturer’s instructions. Protein concentration was quantified by Nanodrop (A280) to normalise the data. Lysates were transferred to a Costar 96-well black clear bottom plate (Thermo Fisher) and substrate working solution was added for 30min. Fluorescence was measured in Glomax plate reader (Promega) with the UV module (365nm excitation and 410–465 emission). Relative caspase levels were normalized to the control (TRE-empty untreated condition).

### Phylogenetic tree assembly

LMX1A/B orthologs were identified using protein-Basic Local Alignment Search Tool (P-BLAST, NCBI). Regions of low compositional complexity were masked to avoid misleading results and an e-value threshold of 10^-80^ was used. Representative organisms from every phylum/family were selected. Protein sequences were aligned using MUSCLE software (EMBL). To generate the LMX1A/B phylogenetic tree, Molecular Evolutionary Genetics Analysis 7.0 (MEGA) software (Pennsylvania) was used (Hall, 2013). A maximum likelihood method with 2000 bootstraps and Jones-Taylor-Thornton (JTT) substitution model was applied—the best-fit substitution model found for our alignments.

### Bioinformatics

Putative LMX1A/B sites in candidate promoter sequences were predicted using MatInspector (Genomatix) (Cartharius, Frech et al., 2005), which identifies transcription factor binding sites using frequency matrices based on published experimental data. Each matrix has an associated random expectation value, indicating how well a matrix is defined (being 0.08% and 0.01% for LMX1A and LMX1B, respectively). This process assigns a maximum score (i.e. probability index), and sequences with relative scores >80%—a commonly used threshold for TFBS (computational framework for transcription factor binding site) analyses using PSSMs (Position Specific Scoring Matrix). To identify putative LMX1A/B LC3-interacting regions (LIR motifs), we used the iLIR Autophagy database (https://ilir.warwick.ac.uk) (Jacomin, Samavedam et al., 2016), applying ANCHOR software to predict flanking, stabilisation regions predicted to stabilize the putative binding. To identify the NLS sequence in LMX1A and LMX1B, we used the cNLS Mapper (Kosugi, Hasebe et al., 2009) which predicted the NLS sequence with a score of 7 and 10, respectively.

### Bioinformatic modelling LC3-LMX1B interaction

Molecular graphics manipulations and visualisations were performed using VMD-1.9.1 and Chimera-1.10.2 [DOI: 10.1002/jcc.20084]. Chimera was used to visualise the 5d94.pdb crystal structure of the LC3B protein and the FYCO1 peptide (Edwards, Rice et al.), and the FYCO1 peptide was used as the “template” to guide the positioning of the LMX1B LIR and flanking sequences. Ultimately the phenylalanine (of FYCO1 LIR) and the required tyrosine (of LMX1B LIR) were aligned and flanking residues on the FYCO1 peptide were altered to mimic the LMX1B residues. Pdb2gmx was used to prepare the assemblies using the v-site hydrogen option to allow a 5fs time step. Hydrogen atoms were added consistent with pH7 and parameterised with the AMBER-99SB-ildn forcefield. Each complex was surrounded by a box 2nm larger than the polypeptide in each dimension, and filled with TIP3P water. The GROMACS-5.1.5 suite of software was used to set up, energy minimise and perform the molecular dynamics (MD) simulations of the resulting assemblies. Random water molecules were replaced by sodium and chloride ions to give a neutral (uncharged overall) box and an ionic strength of 0.15M. Each assembly was subjected to 5000 steps of energy minimisation, velocities were generated with all bonds restrained, prior to molecular dynamics simulations. The LMX1B-LIR/LC3B complex was MD simulated for 75ns, throughout which the LIR peptide remained in contact—this was considered a reasonable binding pose. All simulations were performed as NPT ensembles at 298K using periodic boundary conditions. Short range electrostatic and van der Waals interactions were truncated at 1.4nm while long range electrostatics were treated with the particle-mesh Ewald’s method and a long-range dispersion correction applied. Pressure was controlled by the Berendsen barostat and temperature by the V-rescale thermostat. The simulations were integrated with a leap-frog algorithm over a 5fs time step, constraining bond vibrations with the P-LINCS method and SETTLE for water. Structures were saved every 0.1ns for analysis and each run over 50ns. Simulation data were accumulated on Bristol University Bluecrystal phase 4 HPC. In each case Root Mean Square Deviation (RMSD) was calculated from the trajectories to give an indication of peptide flexibility over the course of the simulations. Root Mean Square Fluctuations (RMSF) were calculated to indicate the flexibility of individual peptide residues over the trajectories. Images were produced with Chimera and Microsoft Paintshop or GIMP (GNU Image Manipulation Program).

### Image analysis and statistics

Fluorescence intensity and puncta quantification were carried out using Fiji. Graphical results were analyzed with GraphPad Prism 7 (GraftPad Software, San Diego, CA), using an unpaired Student’s t-test or one-way ANOVA, as indicated. *p<0.05, **p<0.01 and ***p<0.001. Results are expressed as mean ± SEM or mean ± SD, as indicated.

## ACKNOWLEDGEMENTS

We are grateful to the Wolfson Bioimaging facility for confocal microscopy and Incucyte support, and in particular Stephen Cross for assistance with the neurite growth analysis assay. This work was supported by a Wellcome Trust Ph.D. studentship awarded to N.J-M. through the Dynamic Cell Biology programme; grant number 083474. P.S. was supported through a Medical Research Council DTG studentship. Z.A. was supported by the BBSRC and EPSRC through the BrisSynBio Synthetic Biology Research Centre (BB/L01386X1).

## AUTHOR CONTRIBUTIONS

J.D.L. and N.J-M. conceived and designed the project. J.D.L. supervised the project and analysed data. N.J-M. carried out all wet experiments. D.K.S. and R.B.S. performed the computational modelling. P.S. and M.C. contributed to the mDAN differentiation protocol and P.S. provided mDAN cell culture methodology instruction. R.W. provided reagents and guidance on LMX1B biology. Z.A. carried out the Seahorse bioenergetics experiments. J.D.L and N.J-M. co-wrote the manuscript, all authors critically reviewed and edited the manuscript.

## DECLARATION OF INTERESTS

The authors declare no competing interests.

---

**Table EV1.**
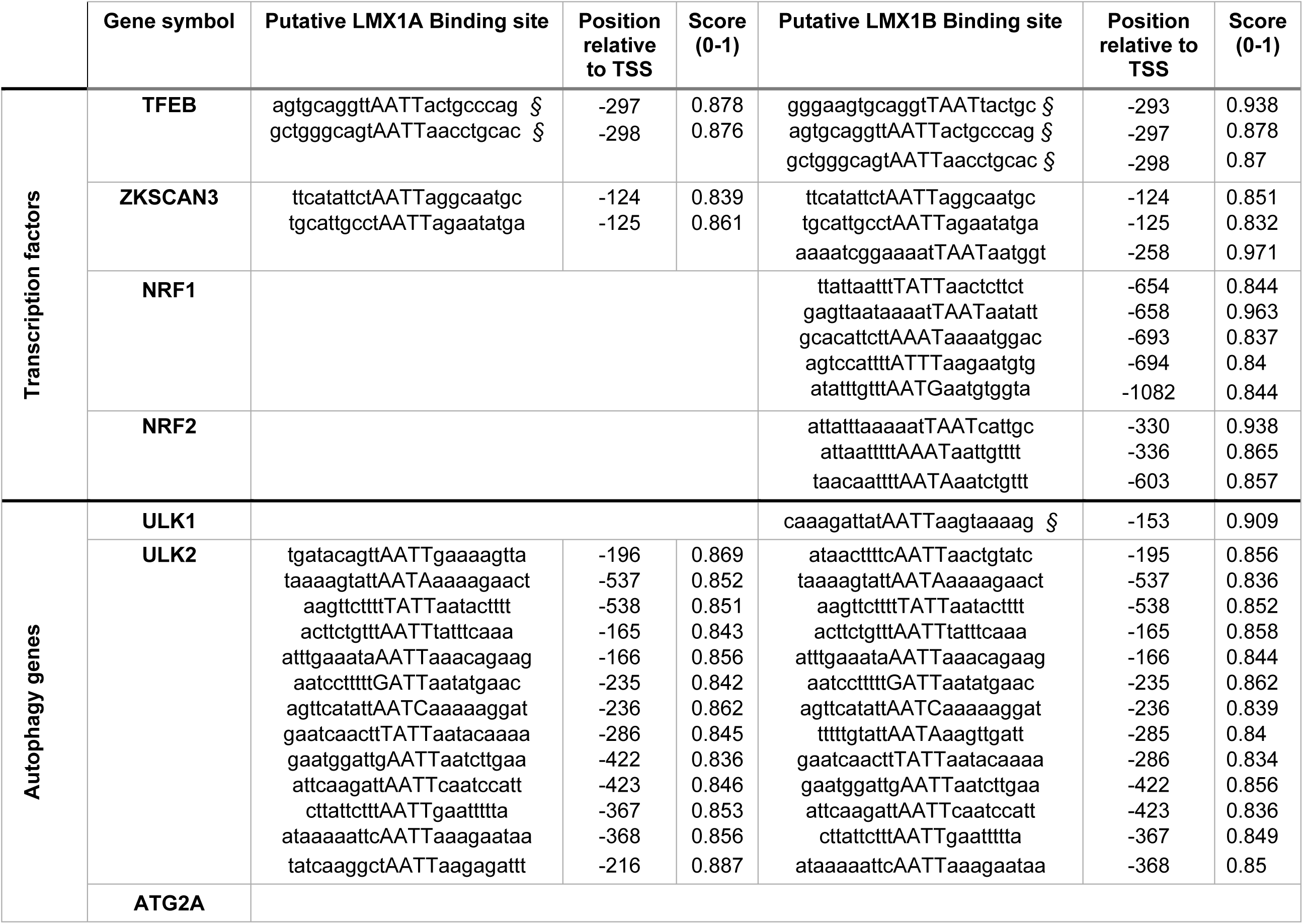
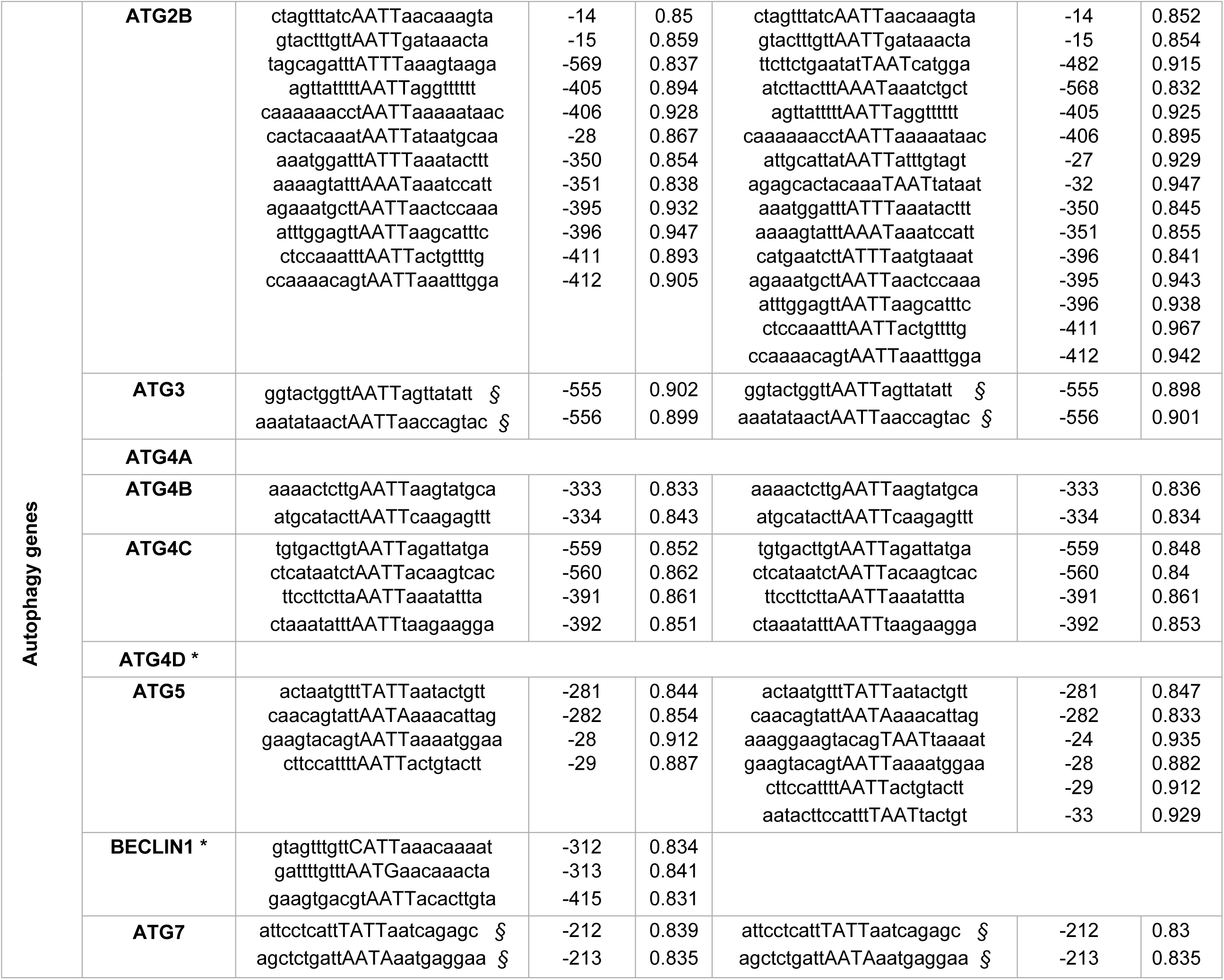
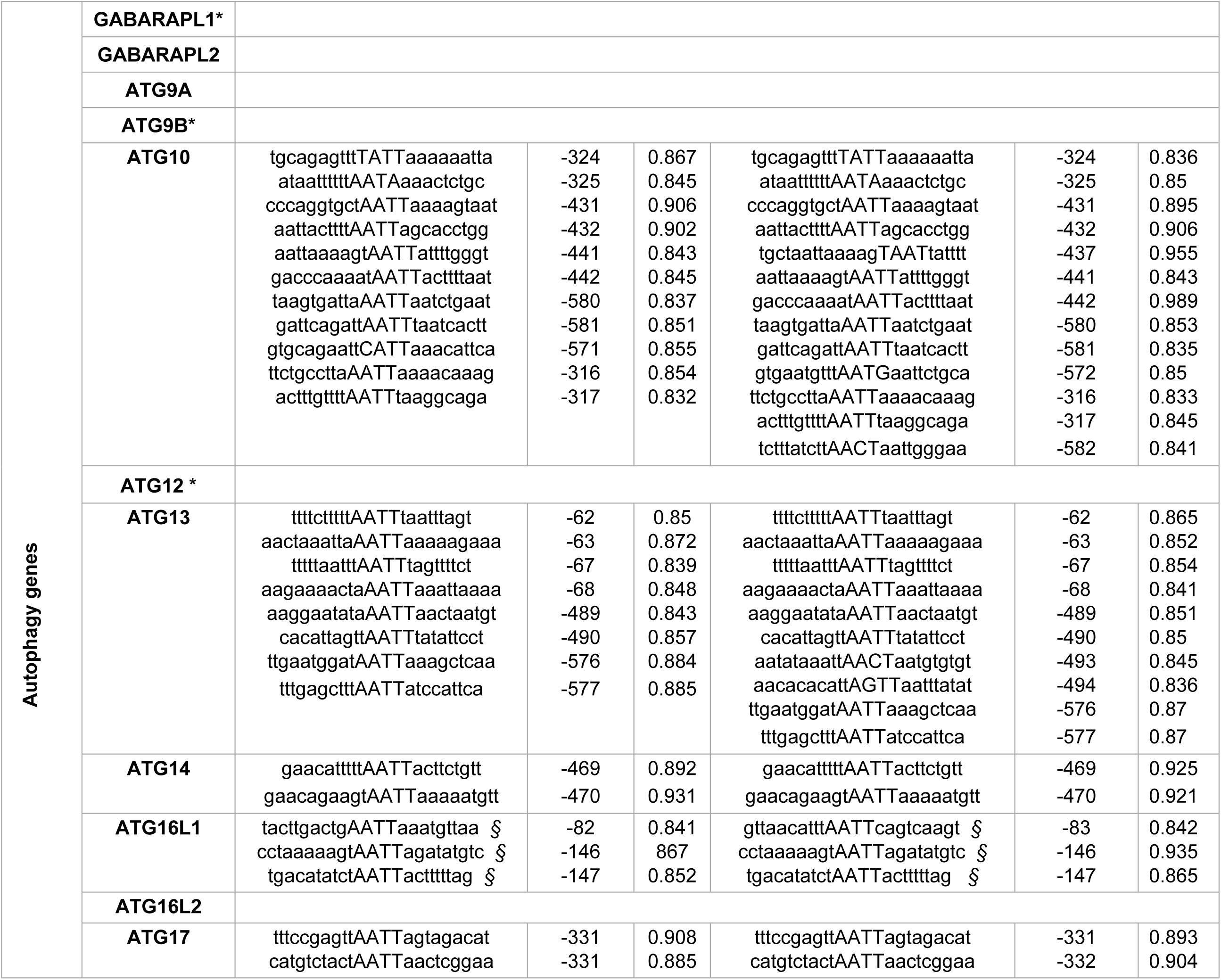
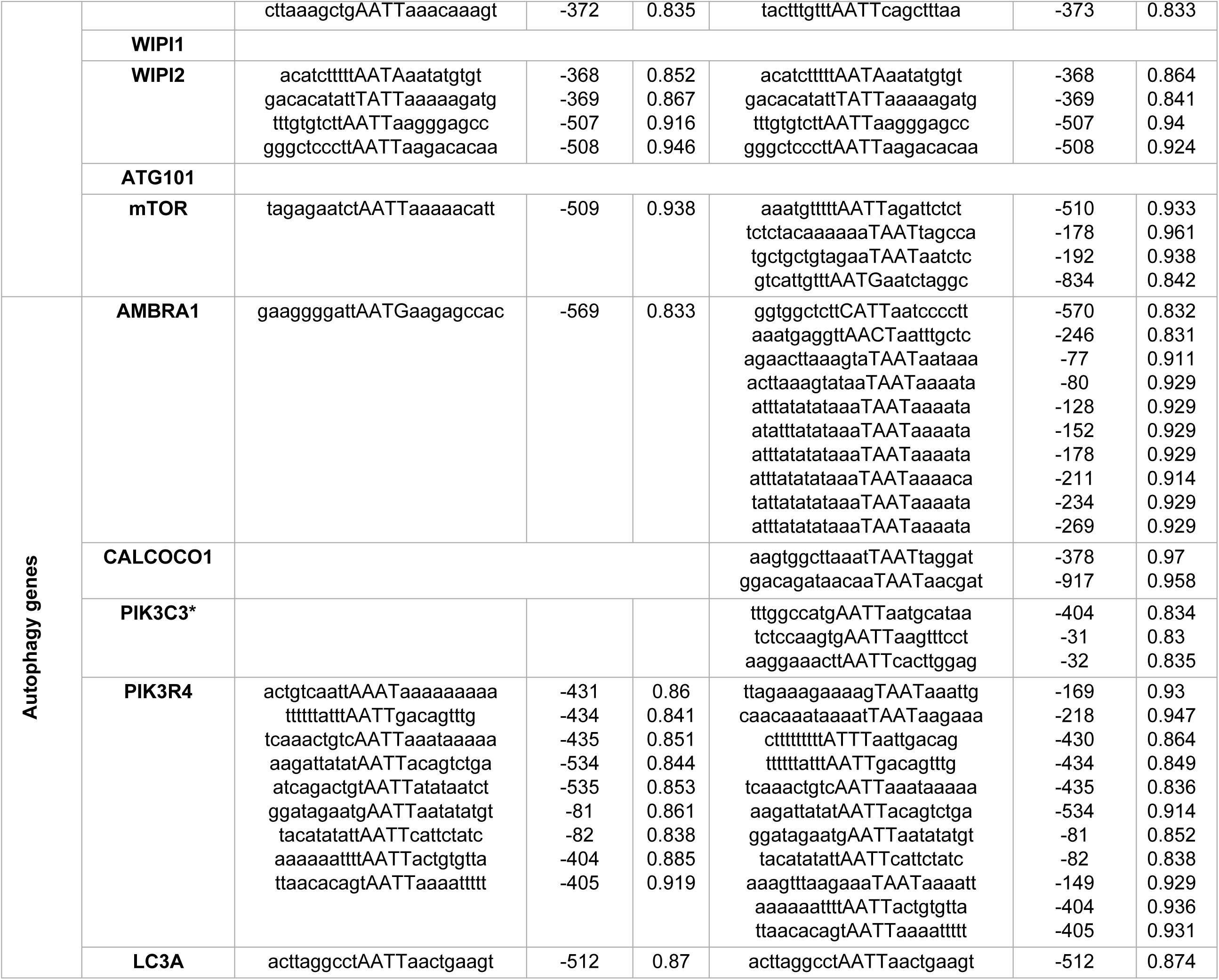
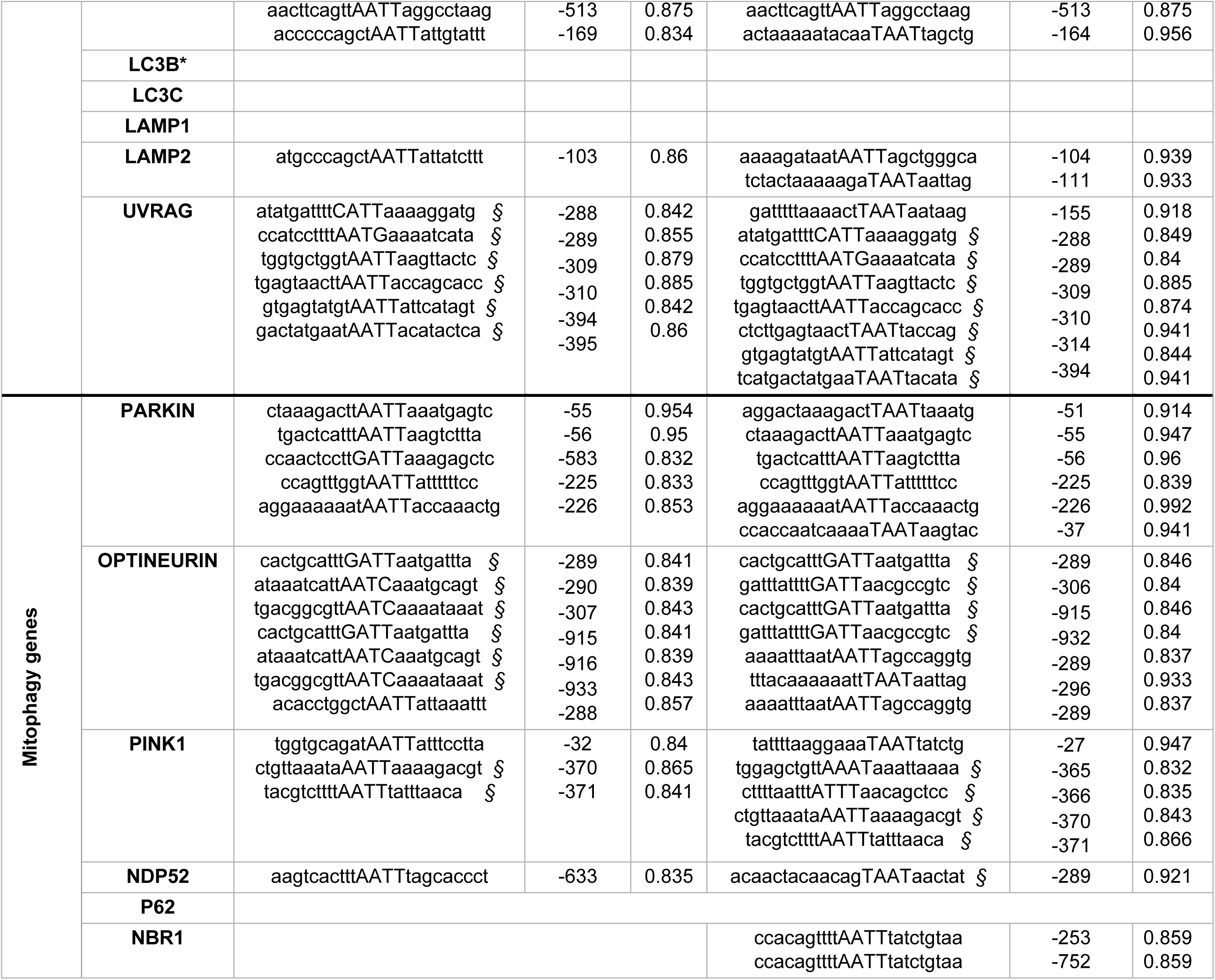
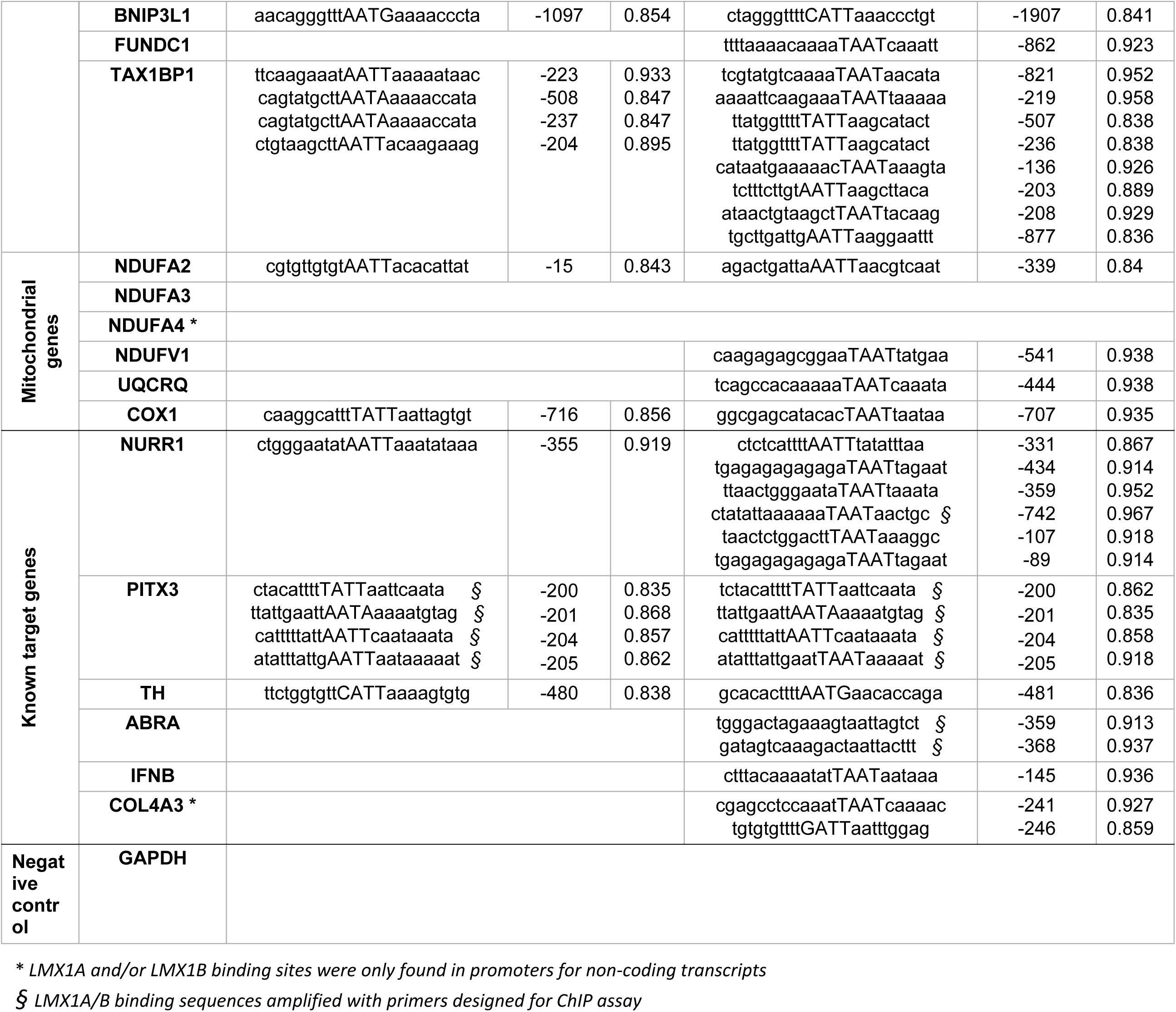
MatInspector promoter FLAT sequence analysis for selected candidate genes.

## EXTENDED VIEW FIGURE LEGENDS

**Figure S1. LMX1A and LMX1B are evolutionary conserved mDAN transcription factors**

(A) Phylogenetic analysis by the Maximum Likelihood method based on the JTT matrix-based model (Bergman et al., 2009). Initial tree for the heuristic search was obtained automatically by applying Neighbor-Join and BioNJ algorithms to a matrix of pairwise distances estimated using a JTT model, and then selecting the topology with superior log likelihood value. A discrete Gamma distribution was used to model evolutionary rate differences among sites (5 categories (+G, parameter = 0.6637)). The analysis involved 23 amino acid sequences. All positions containing gaps and missing data were eliminated. There was a total of 186 positions in the final dataset.

(B) Bootstrap consensus tree inferred from 2000 replicates (Felsenstein, 1985). Only branches that appeared in more than 50% bootstrap replicates are shown. The percentage of replicate trees in which the associated taxa clustered together in the bootstrap test (2000 replicates) are shown next to the branches (Jones, Taylor et al., 1992). Evolutionary analyses were conducted in MEGA7 (Kumar, Stecher et al., 2016).

(C) LMX1A/LMX1B form heterodimers. GFP-TRAP immunoprecipitation of stable TRE-mCherry-LMX1B HEK293T cells previously transfected with either TRE-GFP, TRE-GFP-LMX1A or TRE-LMX1B. Expression was induced by doxycycline and samples were blotted for GFP and mCherry.

(D) Rescue of LMX1B suppression in HEK293T cells were double co-transfected with 50nM Smartpool LMX1B siRNA and codon optimized, siRNA-resistant LMX1B plasmid (or empty vector as a control). Values are mean ± SD of triplicate samples from a single knockdown/rescue experiment, representative of 2 experiments with similar results, and were normalised to GL2. Student’s t test: *p<0.05 and ** p<0.01; siRNA + empty vector vs. siRNA + LMX1B.

**Figure S2. Human mDAN responses during LMX1A/LMX1B suppression and overexpression**

(A, B) Incucyte analysis of neurite length over 4 days in iPSC-derived mDANs transduced with hsyn-GFP/mcherry-U6-shRNA lentiviruses as indicated at (A) D>35 and (B) D<17. mDANs were virally transduced on the day of plating. Shown are means from 3 wells of a single representative experiment. For clarity, error bars have been omitted.

(C) qRT-PCR analysis of mRNA levels following shRNA suppression of LMX1A or LMX1B in young (D17) cultures. Significant reductions in TH, TUJ1, NURR1 and MSX1 expression observed following LMX1A shRNA. LMX1B suppression increases LMX1A levels. Student’s t test: *p<0.05, ** p<0.01 and *** p<0.001.

(D) LMX1A/LMX1B overexpression in iPSC-derived mDANs and autophagy gene expression. Schematic representation of the hsyn promoter modified Tet-On system (top). TRE-GFP-LMX1A and/or TRE-GFP-LMX1B overexpression (or TRE-GFP only as a control) via the Tet-On system in D30 iPSC-derived mDANs (bottom). Expression was induced with the addition of doxycycline (500ng/mL) for 2 days. mRNA levels of the indicated genes were determined by qRT-PCR and normalized by GAPDH levels. Data show means ± SD of triplicate wells from a single representative experiment. Student’s t test: *p<0.05, ** p<0.01 and *** p<0.001 vs. TRE-GFP control.

**Figure S3. LMX1A binds ATG8 family members in vitro**

(A) Domain schematic of human LMX1A and LMX1B showing the position of a possible LIR motif in LMX1A (left). Alignment of the putative LMX1A LIR in different species (right).

(B) GFP-TRAP co-precipitation of LMX1A with GFP-ATG8 family members in HEK293T cells. 5% protein lysate from equivalent GFP-expressing cells is shown as “input”. Arrow indicates the position of the LMX1A band.

(C) GFP-TRAP co-precipitation of LMX1A with wild-type of LIR docking mutant (F52A/L53A) GFP-LC3B in HEK293T cells. 5% of protein lysates were used as control for protein expression (inputs). Arrow indicates position of the LMX1A band.

(D) GFP-TRAP pull-downs in lysates of HEK293T cells expressing LIR mutant (Y290A/L293A) LMX1A and GFP-LC3B. Arrow indicates position of the LMX1A band.

(E) GFP-TRAP immunoprecipitation of nuclear and cytosolic fractions from lysates of HEK293T cells co-expressing GFP-LC3A, B, C and LMX1A under basal conditions. Arrow indicates position of the LMX1A band.

(F) GFP-TRAP immunoprecipitation of nuclear and cytosolic fractions from lysates of HEK293T cells co-expressing GFP-LC3B and LMX1A. Comparisons of pull-downs in full nutrients following 2h starvation. Arrow indicates position of the LMX1A band.

(G) LMX1A turnover in HEK293T cells. HEK293T cells were transfected with LMX1A and treated with cycloheximide in the absence or presence of BafA1 or MG132.

(H) GFP-TRAP immunoprecipitation of LMX1A in HEK293T cells co-expressing wild-type, acetylation-deficient (K49R/K51R) and acetylation mimic (K49Q/K51Q) GFP-LC3B. Arrow indicates position of the LMX1A band.

**Figure S4. In silico modelling of the LMX1B-LC3B interaction**

(A) Frames from a computer simulation of the LIR-dependent LMX1B-LC3B interaction (see **Video S1**) showing the dynamics of the putative LMX1B LIR motif docked by molecular replacement in the position adopted by the FYCO1 LIR (5d94.pdb). LC3B is shown in cyan; LMX1B LIR in magenta.

(B) Comparison of the final LMX1B LIR pose and the position of the N-terminal ATG4B LIR identified in the ATG4B/LC3B crystal lattice (Satoo et al., 2009).

**Figure S5. LMX1B binding to ATG8 proteins is restricted to the nucleus**

HEK293T cells expressing GFP-ATG8 proteins and LMX1B-FLAG were separated into nuclear and cytoplasmic fractions which were subjected to GFP-TRAP.

**Movie EV1. Computer simulation of the interaction between the LMX1B LIR and LC3B**

Movie of an example computer simulation of the LIR-dependent LMX1B-LC3B interaction (relates to **Figure EV3C**). The LMX1B LIR motif was docked by molecular replacement in the position adopted by the FYCO1 LIR (5d94.pdb). LC3B is shown in cyan; LMX1B LIR in magenta. The movie spans 50ns.

